# Complement activation by IgG subclasses is governed by their ability to oligomerize upon antigen binding

**DOI:** 10.1101/2024.03.26.586731

**Authors:** Nikolaus Frischauf, Jürgen Strasser, Ellen G.F. Borg, Aran F. Labrijn, Frank J. Beurskens, Johannes Preiner

## Abstract

Complement activation through antibody-antigen complexes is crucial in various pathophysiological processes such as infections, inflammation, and autoimmunity, but is also utilized in immunotherapies to eliminate infectious agents, regulatory immune cells, or cancer cells. Although the tertiary structures of the four IgG antibody subclasses are largely identical, complement recruitment and further activation depend strongly on subclass, which is commonly explained by the respective affinity for C1, the first component of the classical complement pathway. Contradicting this established view, we here demonstrate that complement activation by different IgG subclasses is determined by their varying ability to form IgG oligomers on antigenic surfaces large enough to multivalently bind and activate C1. We directly visualize the resulting IgG oligomer structures and characterize their distribution by means of high-speed atomic force microscopy (HS-AFM), quantify their complement recruitment efficiency from quartz crystal microbalance (QCM) experiments, and characterize their ability to activate complement on tumor cell lines as well as in vesicle-based complement lysis assays. We present a mechanistic model of the multivalent interactions that govern C1 binding to IgG oligomers and use this model to extract affinities and kinetic rate constants from real-time interaction QCM data. Together, our detailed characterization yields a comprehensive view on the parameters that govern complement activation by the different IgG subclasses, which may inform the design of future antibody therapies.

## Introduction

Immunoglobulin G (IgG) is the most abundant antibody isotype in human serum and a highly potent effector molecule. IgG is divided into four subclasses designated IgG1, IgG2, IgG3, and IgG4 in order of decreasing abundance. Clearance of IgG-opsonized pathogens or cells is facilitated by various effector mechanisms of the innate immune system, including the classical complement pathway, an amplifiable cascade of soluble zymogens abundant in blood and other extracellular fluids. It is initiated by the binding of zymogen C1 (C1qr_2_s_2_), consisting of the hexavalent recognition protein C1q (made of six collagen arms assembled in a “bunch of tulips” like structure, each ending in a globular IgG-Fc binding domain; gC1q) and a heterotetramer of serin proteases C1r and C1s (C1qr_2_s_2_), to antibody-antigen complexes (1–3). Rearrangements within C1 as a consequence of antibody recognition by C1q activates the proteases that are then able to cleave complement proteins C4 and C2 into C4a,b and C2a,b with the respective b products potentially being covalently deposited on the antigenic membrane. In this way, the formation of downstream enzymatic complexes, the C3 and C5 convertases (C4b2b and C4b2b3b) is induced, which then may lead to the formation of the membrane attack complex (MAC, C5b-9), a lytic pore, that is inserted into the target cell membrane.

Although the tertiary structures of the four IgG subclasses are largely comparable, with the most prominent difference being the Fab/Fc connecting hinge region (length of 15 aa, 12 aa, 62aa, and 12 aa in IgG1-4, respectively), complement recruitment and further activation depend strongly on subclass, which is commonly attributed to the respective affinity for C1q (4) decreasing in the order IgG3>IgG1>IgG2>IgG4 (5). However, since the corresponding equilibrium dissociation constants of the monovalent interactions range between 34 µM (IgG3) and 229 µM (IgG4), significant C1 binding at physiological concentrations (∼0.17 µM (6)) requires multivalent interactions between C1 and multiple IgGs, irrespective of subclass. We have recently shown that antigen-mediated oligomerization of IgG1 via non-covalent Fc-Fc interactions is crucial for engaging the complement system through C1(q) binding. While antigen bound IgG1 may exist as a distribution of oligomers ranging from monomers to hexamers, only tetramers, pentamers, and hexamers were capable of inducing complement dependent cytotoxicity (CDC), as a minimum of four gC1q headpieces was required to bind in close proximity to adjacent subunits within IgG1 oligomers (7). Accordingly, tetravalent C1 binding to non-adjacent subunits within an IgG1 hexamer did not induce CDC, underlining the necessity for a particular binding geometry and valency that results in sufficient compaction of C1q arms to induce conformational rearrangements that allow C1r to activate C1s (2). While the formation of IgG oligomers has only directly been shown for the IgG1 subclass (1, 2, 7, 8) and more recently for IgG3 (9), there are indications that IgG Fc point mutations that enhance oligomerization and CDC efficacy of IgG1 against a certain target, also affect its IgG2, IgG3, and IgG4 variants in a similar manner (1, 10). It is thus likely that all IgG subclasses need to assemble into similar oligomeric structures that result in a common Fc-platform that allows for at least tetravalent C1 binding to adjacent IgG monomers within the oligomer to activate complement.

In this study, we demonstrate that IgG oligomerization is a general mechanism inherent to all IgG subclasses and the main pre-requisite for complement C1 binding and activation. We performed CDC assays to assess and compare the efficacy of IgG subclasses as well as the effect of an IgG1-oligomerization enhancing Fc point mutantion on all IgG subclasses on killing different lymphoma cell lines. Using high-speed atomic force microscopy (HS-AFM), we visualize the respective oligomer structures formed on antigenic supported lipid bilayers and characterize the oligomerization propensity of IgG subclasses as well as CDC-enhanced Fc point mutants thereof. We further study complement C1 and C1q binding to IgG1-4 oligomers on antigenic membranes in quartz crystal microbalance (QCM) experiments to determine kinetic rate constants, equilibrium dissociation constants and resulting complement recruitment efficiencies at various antigen densities. Finally, we perform liposomal vesicle-based complement lysis assays to link our single molecule observations to terminal complement activation on antigen-coated vesicles. We highlight a complex interplay of innate antibody properties and multivalent IgG-C1(q) interactions, that determine the IgG subclass specific differences in complement recruitment and activation.

## Results

### CDC efficacy depends on IgG subclass and can be enhanced by the E430G point mutation

First, we assessed the efficacy of IgG1-4 and corresponding E430G Fc point mutants, which increased oligomerization and CDC of IgG1 (7, 8, 11), in CDC assays involving different lymphoma cell lines (Fig. 1, Table 1). CDC of human Burkitt’s lymphoma RAJI and DAUDI cells by human anti-CD20 IgG-7D8 was comparably efficient for IgG1-7D8 (Fig. 1A) and IgG3-7D8 (Fig. 1C), whereas no CDC was induced by IgG2-7D8 (Fig. 1B) and was only detected at highest IgG concentrations in case of IgG4-7D8 not reaching the maximum value of IgG1-7D8 (Fig. 1D). Application of anti-CD20 IgG-7D8 subclass variants to WIEN-133 cells yielded vastly comparable EC50 values for CDC albeit maximum lysis was markedly reduced for IgG3-7D8 and IgG4-7D8 to ∼ 20% as compared to their IgG1 variant. Introduction of the E430G mutation into the anti-CD20 IgGs resulted in significantly lowered EC50 values for lysis by IgG1-7D8-E430G in all three cell lines, and strongly enhanced CDC induced by IgG2-7D8-E430G and IgG4-7D8-E430G, both reaching the EC50 values of IgG1-7D8 and IgG3-7D8. IgG3-7D8-E430G, on the other hand, exhibits only modestly improved EC50 values for CDC of RAJI and DAUDI cells compared to IgG3-7D8, and an improved maximum lysis (40% of the IgG1-7D8) on WIEN-133 cells at a similar EC50 value as IgG3-7D8. Applying human anti-CD52 IgG1-CAMPATH and IgG3-CAMPATH to WIEN-133 (human Burkitt’s lymphoma) cells induced CDC at higher EC50 values as compared to the respective anti-CD20 antibodies, and IgG3-CAMPATH reached only 30% of the maximum lysis of IgG1-CAMPATH. Introduction of the E430G mutation into anti-CD52 IgGs resulted in a potent increase of CDC by IgG1-CAMPATH-E430G (∼35-fold lowered EC50 and 60 % increased maximum lysis) and a significant improvement of CDC by IgG3-CAMPATH-E430G (∼3-fold lowered EC50 and maximum lysis level similar to IgG1-CAMPATH). CDC thus depended on the subclass, with IgG1 being the most effective, closely followed by IgG3 in all cell lines tested, while IgG2 and IgG4 hardly induced CDC. The E430G point mutation significantly lowered EC50 values for CDC throughout all tested cell lines and IgG subclasses, suggesting that comparable to its effect on IgG1 oligomerization, it increases oligomerization of IgG2-4.

**Figure 1.**
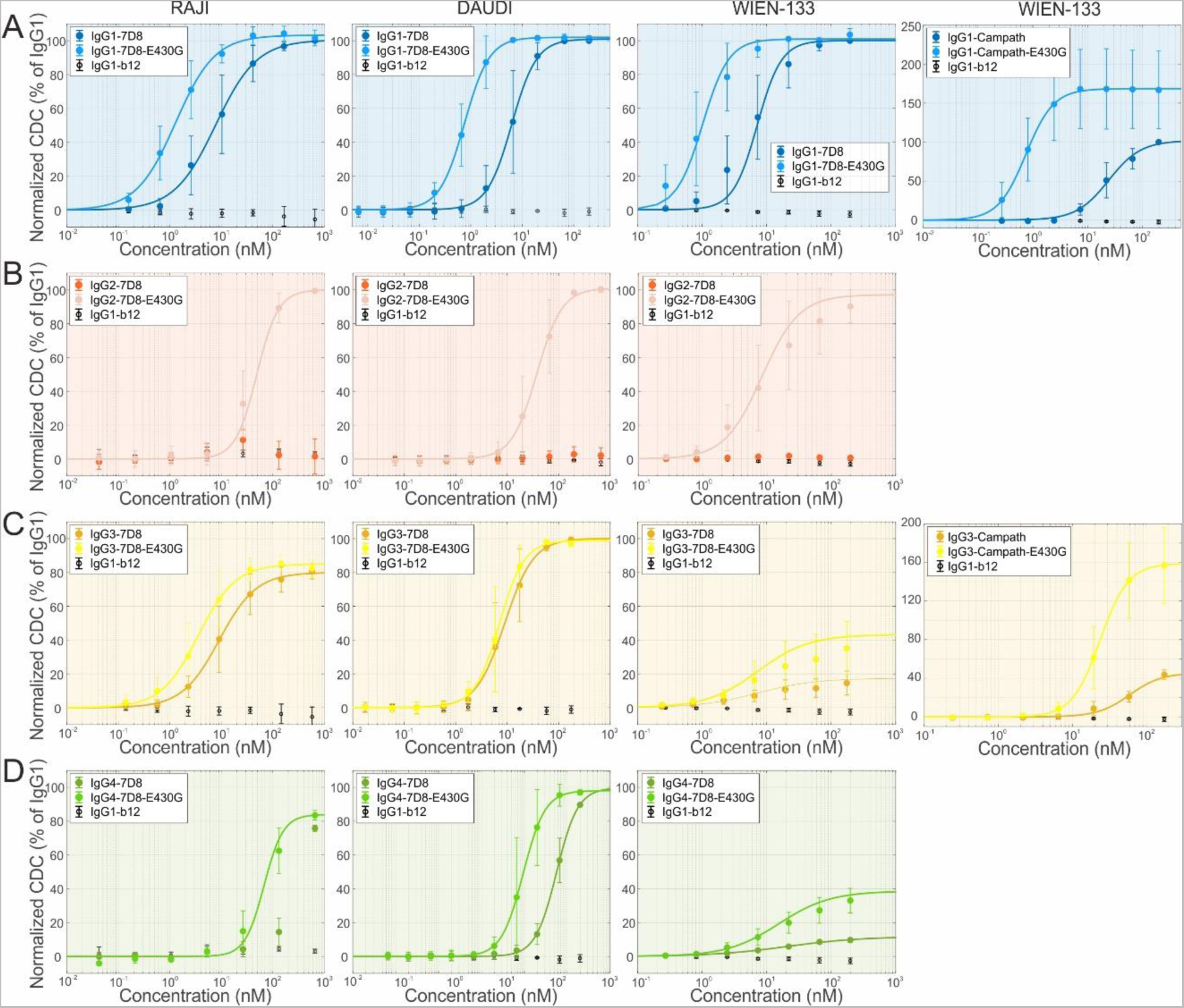
Complement activation by different IgG subclass variants on different lymphoma cell lines and antigenic targets. **(A)** CDC of anti CD20 antibodies IgG1-7D8 and IgG1-7D8-E430G applied to RAJI, DAUDI, and WIEN-133 cells (panel 1-3). Anti CD52 antibodies IgG1-Campath and IgG1-Campath-E430G applied to WIEN-133 cells (panel 4). **B)** CDC of anti CD20 antibodies IgG2-7D8 and IgG2-7D8-E430G applied to RAJI, DAUDI, and WIEN-133 cells. **C)** CDC of anti CD20 antibodies IgG3-7D8 and IgG3-7D8-E430G applied to RAJI, DAUDI, and WIEN-133 cells (panel 1-3). Anti CD52 antibodies IgG3-Campath and IgG3-Campath-E430G applied to WIEN-133 cells (panel 4) **D)** CDC of anti CD20 antibodies IgG4-7D8 and IgG4-7D8-E430G applied to RAJI, DAUDI, and WIEN-133 cells. Solid lines are fits of dose-response curves to the data, EC50 values are displayed in Table 1.

**Table 1.** EC50 (antibody concentration inducing half-maximal lysis) values for CDC of antibody-opsonized cells obtained from fitting a dose-response curve of the type 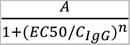 to the CDC data, Fig. 1. Values in brackets are 95% confidence intervals.

| **EC50 (nM)** | **RAJI** | **DAUDI** | **WIEN** |
| --- | --- | --- | --- |
| IgG1-7D8 | **5.6** (4.1, 7.0) n=9 | **6.3** (4.7, 7.9) n=3 | **7.2** (6.3, 8.1) n=3 |
| IgG1-7D8-E430G | **1.0** (0.8, 1.2) n=9 | **0.8** (0.6, 0.9) n=3 | **1.0** (0.9, 1.2) n=3 |
| IgG1-Campath |  |  | **24.2** (16.9, 31.5) n=5 |
| IgG1-Campath-E430G |  |  | **0.7** (0.4, 1.1) n=5 |
| IgG2-7D8 | **n.s.** | **n.s.** | **n.s.** |
| IgG2-7D8-E430G | **4.9** (3.7, 6.1) n=3 | **3.8** (2.8, 4.7) n=3 | **8.3** (6.1, 10.5) n=3 |
| IgG3-7D8 | **5.6** (4.0, 7.2) n=9 | **9.0** (6.1, 11.8) n=3 | **7.3** (1.1, 13.5) n=3 |
| IgG3-7D8-E430G | **3.2** (2.5, 3.9) n=9 | **7.0** (5.1, 8.9) n=3 | **7.3** (3.0, 11.7) n=3 |
| IgG3-Campath |  |  | **59.6** (48.2, 71.0) n=3 |
| IgG3-Campath-E430G |  |  | **23.7** (15.3, 32.1) n=3 |
| IgG4-7D8 | **n.s.** | **56.4** (44.4, 68.5) n=3 | **20.9** (2.8, 38.9) n=3 |
| IgG4-7D8-E430G | **6.8** (5.4, 8.2) n=3 | **9.4** (6.1, 12.7) n=3 | **15.5** (8.2, 22.7) n=3 |

### Antigen dependent oligomerization naturally occurs in all IgG subclasses

Given that the E430G point mutation that increased oligomerization and CDC of IgG1 (7, 8, 11) similarly leads to increased CDC of IgG2-4 E430G variants, we hypothesize that IgG oligomerization is inherent to all subclasses and governs their ability to stably bind and activate C1. We thus prepared supported lipid bilayers (SLB) containing 5% dinitrophenyl (DNP)-labeled lipids (7, 8) to characterize the ability of different IgG subclass variants and mutants to form IgG oligomer populations on antigenic membranes in HS-AFM experiments. While incubation with 33 nM DNP-unspecific IgGs for 5 min did not result in any binding, incubation with IgG1-DNP, IgG2-DNP, IgG3-DNP, and IgG4-DNP under similar conditions generated sparse distributions of differently sized assemblies on top of the SLBs, characteristic for IgG oligomer distributions as previously found for IgG1 (7, 8). Unlike the other subclasses, addition of IgG2 frequently lead to detachment of the SLBs from the mica support so that only smaller IgG2-bearing lipid patches that remained firmly attached to the support could be analyzed. We mainly observed isolated, likely bivalently bound, IgGs with heights of 4-7 nm (IgG1, IgG2, and IgG4) and 5-12 nm (IgG3) and smaller fraction of higher-order IgG assemblies with heights ranging up to 11 nm (IgG1, IgG2, and IgG4) and 15 nm (IgG3), respectively (Fig. 2A-D). The E430G single point mutation only had a recognizable effect on the height distribution of IgG1-DNP-E430G with the second height population between 6 and 10 nm becoming more prominent as compared to IgG1-DNP (Fig. 2 E). For the other subclasses, no such change in height distribution was apparent (Fig. 2F-H). The triple mutants IgG1-DNP-RGY, IgG2-DNP-RGY, IgG3-DNP-RGY, and IgG4-DNP-RGY (E345R, E430G, and S440Y) which were previously shown to efficiently associate into IgG hexamers in solution (1, 12, 13) exhibited the largest fraction of large IgG assemblies as evident from the broadening of the height distributions and the appearance of a second, higher peak for all four subclasses (Fig. 2I-L). Interestingly, the IgG3 variants always contained a fraction that protruded further (up to 15 nm) from the antigenic membranes then the respective IgG1, IgG2, and IgG4 variants (up to 10 - 11 nm). They also exhibited a higher resistance to increased HS-AFM imaging forces than the other three IgG subclasses, which could be more easily removed from the membranes when intentionally lowering the HS-AFM setpoint amplitude, potentially indicating bivalent attachment to antigenic epitopes of IgG3 molecules within oligomers.

**Figure 2.**
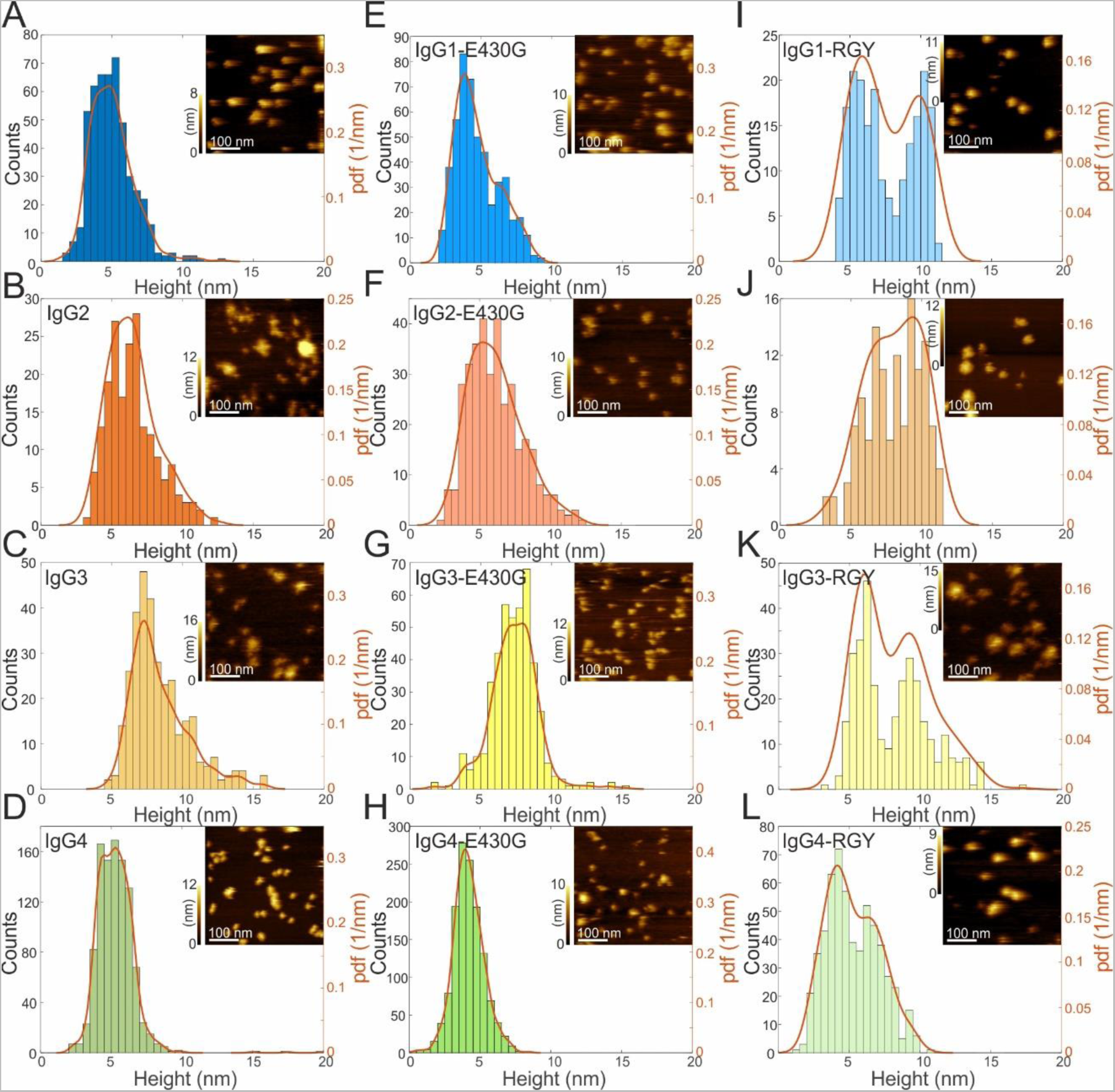
HS-AFM height assessment of different IgG variants bound to antigenic surfaces. **(A, E, I)** Height distributions (histograms and probability density functions) of IgG1-DNP, IgG1-DNP-E430G, and IgG1-DNP-RGY bound to DNP-SLBs. **(B, F, J)** Height distributions of IgG2-DNP, IgG2-DNP-E430G, and IgG2-DNP-RGY bound to DNP-SLBs. **(C, G, K)** Height distributions of IgG3-DNP, IgG3-DNP-E430G, and IgG3-DNP-RGY bound to DNP-SLBs. **(D, H, L)** Height distributions of IgG4-DNP, IgG4-DNP-E430G, and IgG4-DNP-RGY bound to DNP-SLBs. Insets represent typical HS-AFM images of the respective IgG variant bound to DNP-SLBs.

### IgG3 oligomers are structurally distinct from other IgG subclasses

To further examine these topographical differences, we obtained higher resolution images of IgG3-DNP assemblies on antigenic membranes as exemplified in Fig. 3 A, taken from HS-AFM movie S1. Judging from the lateral size, symmetry, and mobility of the respective assembly, we identified them as monomers and hexamers as indicated in Fig. 3A. Tracking these IgG oligomers over time enabled us to evaluate their heights in each image frame, from which we again generated height histograms (Fig. 3B). The IgG3-DNP monomers recorded in this HS-AFM movie were characterized by a height of 5.8 nm that fluctuated by ± 0.7 nm during the experiment, whereas the hexamers protruded on average 13.2 nm from the antigenic membrane and fluctuated by ± 1.3 nm (mean ± s.d. overall image frames and individual particles, respectively). Fig. S1A depicts the first HS-AFM image frame of movie S2, another example of IgG3 oligomers bound to an antigenic membrane, however, additionally to monomers and a hexamer, a tetramer was followed and analyzed over time. While height histograms of monomers and hexamers were consistent with the respective distributions from Fig. 3B, the height distribution of the tetrameric IgG3 was located in between the monomers and the hexamer exhibiting an average height of 9.4 and fluctuations of ± 1.7 nm. Repeating these experiments and height analysis for IgG1 (Fig. 3C, movie S3) resulted in much narrower distributions with average heights of 7.3 and 11.1, and height fluctuations of ± 0.8 and ± 0.6 for IgG1 monomers and hexamers, respectively. These experiments suggest that the height distributions compiled from a large ensemble of antigenic membrane bound IgG assemblies as depicted in Fig. 2 include overlapping contributions from differently sized IgG oligomers with the lower heights stemming mainly from IgG monomers and the larger ones from the IgG hexamers. Zooming further in on IgG3 hexamers (Fig. 3E taken from movie S4 and Fig. S1C) clearly revealed a central platform consisting of six Fc fragments (9) with an overall shape and diameter as previously found for IgG1 hexamers (7), but with smaller portions that laterally protruded below the platform (indicated by arrows) that were not found in case of IgG1 hexamers (Fig. 3F). These smaller parts slowly changed their position around the central platform over time, however, the fact that they could be resolved by the HS-AFM tip suggests that they represent antigen-bound IgG3 Fabs, since unbound Fabs as present in IgG1 hexamers would be subject to much faster diffusional motion far beyond the time-resolution of the HS-AFM (14).

**Figure 3.**
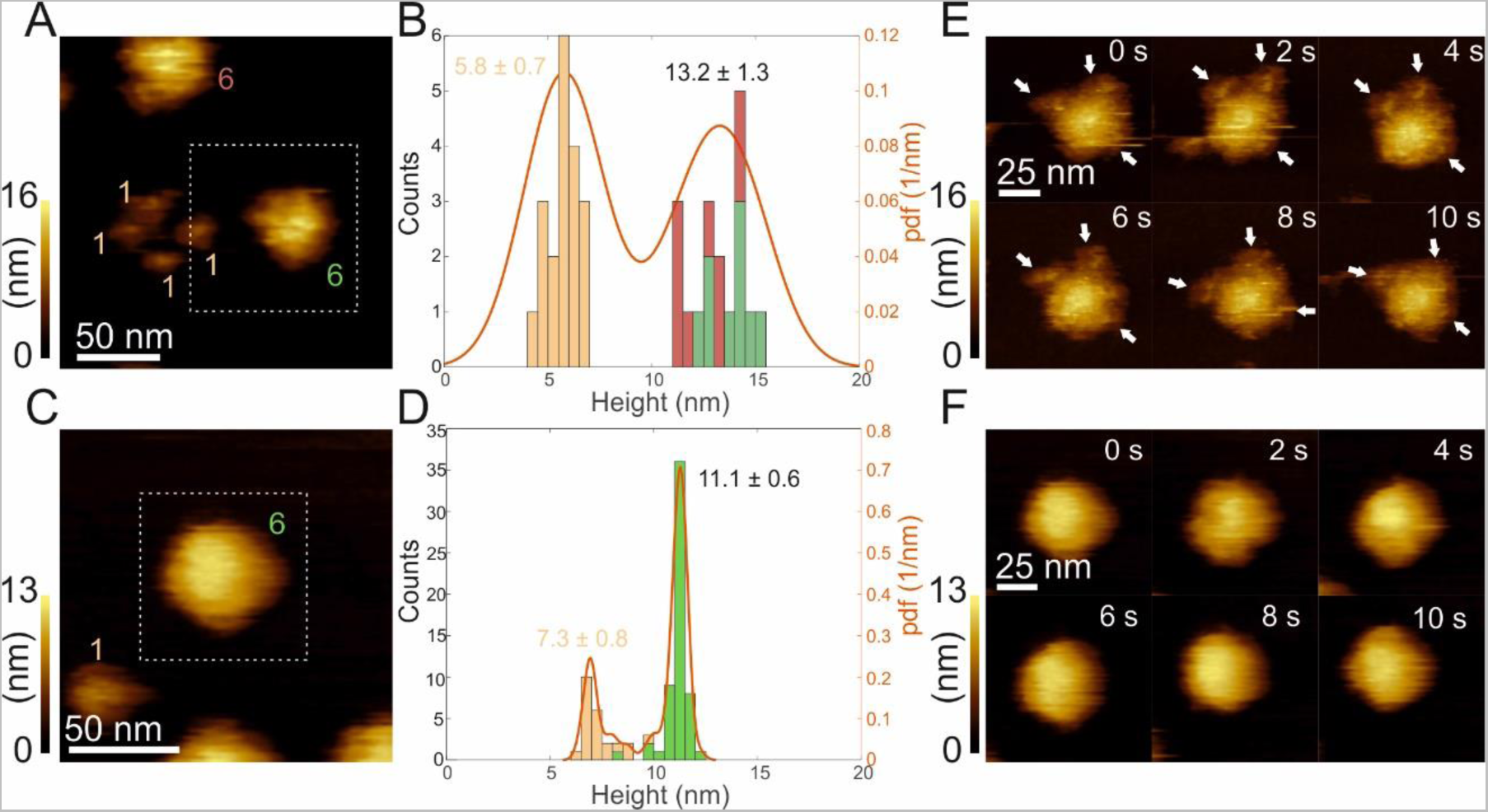
Structural comparison of IgG3 and IgG1 hexamers based on higher-resolution HS-AFM images. **(A)** First frame of HS-AFM movie S1 of two IgG3 hexamers and four IgG3 monomers bound to a DNP-SLB. **(B)** Height histogram generated from height over time recordings of the oligomers in movie S1. Contributions of the respective oligomers are color coded according to (A). Numbers correspond to means ± s.d. over all image frames and individual particles, respectively. **(C)** First frame of HS-AFM movie S3 of an IgG1 hexamer and one IgG1 monomer bound to a DNP-SLB. **(D)** Height histogram generated from the height over time of the oligomers in movie S3. Contributions of the respective oligomers are color coded according to (C). **(E)** High resolution images of an individual IgG3 hexamer (dashed area from (A)) taken from HS-AFM movie S4. Additional smaller structures surrounding the central Fc platform are indicated by arrows. **(F)** High resolution images of an individual IgG1 hexamer (dashed area from (C)).

### Abundance of higher IgG oligomers depends on subclass and increases with E430G point mutation

HS-AFM imaging further allowed us to compile quantitative oligomer distributions for each subclass variant on DNP-SLBs. The molecules were therefore scanned in a non-disrupting manner to gauge their number, height, and shape, followed by an increase in scanning force applied by the HS-AFM cantilever tip to dissociate the oligomers into their constituent individual IgGs (7, 8). Geometric parameters and oligomer decay patterns were combined to assign each IgG assembly its oligomeric state (Fig. 4). Characteristic oligomer distributions were found for all investigated IgGs, with smaller oligomers occurring more frequently than large ones. While IgG1-DNP, IgG2-DNP and IgG3-DNP already formed oligomers up to hexamers under the chosen conditions (Fig. 4A-C), only a few pentamers were detected in the case of IgG4-DNP (Fig. 4D). Introduction of the E430G point mutation increased the oligomerization propensity of IgG1, IgG2 and IgG4 leading to a significantly higher amount of pentamers and hexamers relative to the respective parental IgG, while the abundance of IgG3-DNP-E430G pentamers and hexamers stayed mostly at the same level as IgG3-DNP (Fig. 4E).

**Figure 4.**
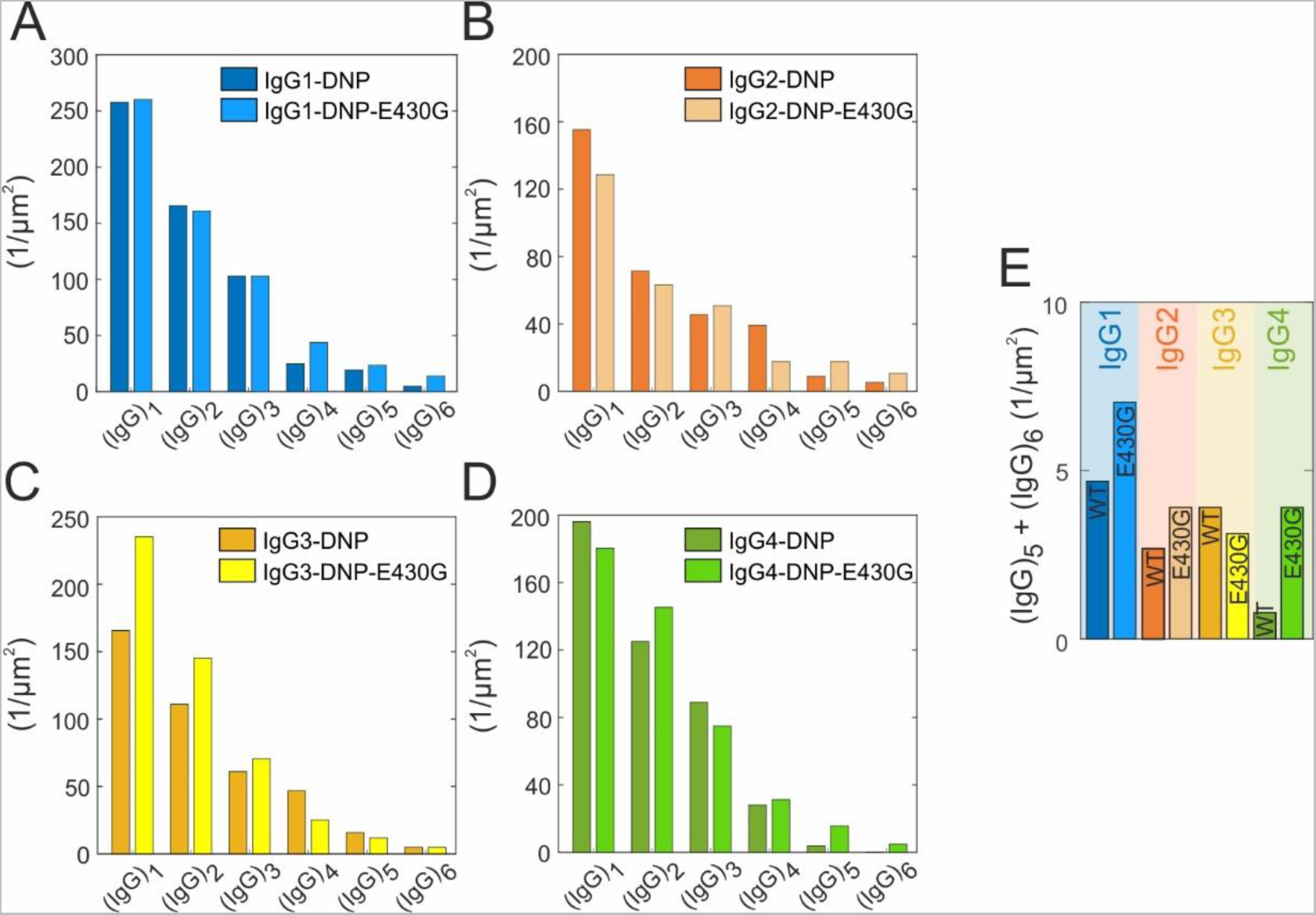
Oligomer distributions of anti-DNP IgG1-4 and the respective E430G point mutants bound to DNP-SLBs examined by HS-AFM. **(A)** IgG1-DNP and IgG1-DNP-E430G. **(B)** IgG2-DNP and IgG2-DNP-E430G. **(C)** IgG3-DNP and IgG3-DNP-E430G. **(D)** IgG4-DNP and IgG4-DNP-E430G. **(E)** Sum of IgG pentamer- and hexamer densities for each IgG variant.

### C1 recruitment efficiency correlates with IgG oligomerization propensity

To address the question of whether the observed oligomerization propensity is directly related to complement C1 fixation, we characterized the ability of the different IgG subclasses and E430G point mutants to recruit C1 to antibody-opsonized antigenic membranes in quartz crystal microbalance (QCM) experiments (8) employing the same DNP-SLBs used in HS-AFM but generated on a QCM senor chip (Fig. 5A). Monitoring the change of oscillation frequency of the DNP-SLB covered SiO_2_-coated quartz crystal, which is proportional to the change in bound mass due to association of molecules out of the constant buffer flow above the chip surface yields characteristic binding curves for lipids, IgGs and subsequently added complement proteins (Fig. S2). We prepared three DNP-SLBs differing in antigen densities (0.1, 0.5, and 5 % DNP) to which the respective IgG variant was allowed to associate for a defined timespan. After establishing a certain IgG density (∼2, 3, and 7 x 10^3^ IgGs/µm^2^, respectively) and removal of solution phase IgGs, C1 was added to the running buffer and the binding signal was followed over time until the end of the dissociation phase in which C1 was again removed from the running buffer (Fig. S3A-B). To better compare the respective C1 binding levels between different experiments, we introduced the ratio of bound IgGs to C1 molecules evaluated at the maximum (max.) C1 level and the final C1 level at the end of the experiment (EOX), respectively (Fig. 5B). Importantly, these ratios of average IgGs needed to recruit a single C1 molecule to our antigenic membranes define the density of IgGs needed to generate a binding site for a single C1, however, they are not necessarily reflecting the number of IgGs actually interacting with a C1 molecule (limited by max. six gC1q binding heads per C1). The comparison of these ratios resulting from max. C1 and EOX C1 enables evaluation of the stability of the established IgG-C1 complexes and its variation with subclass and mutant variants.

**Figure 5.**
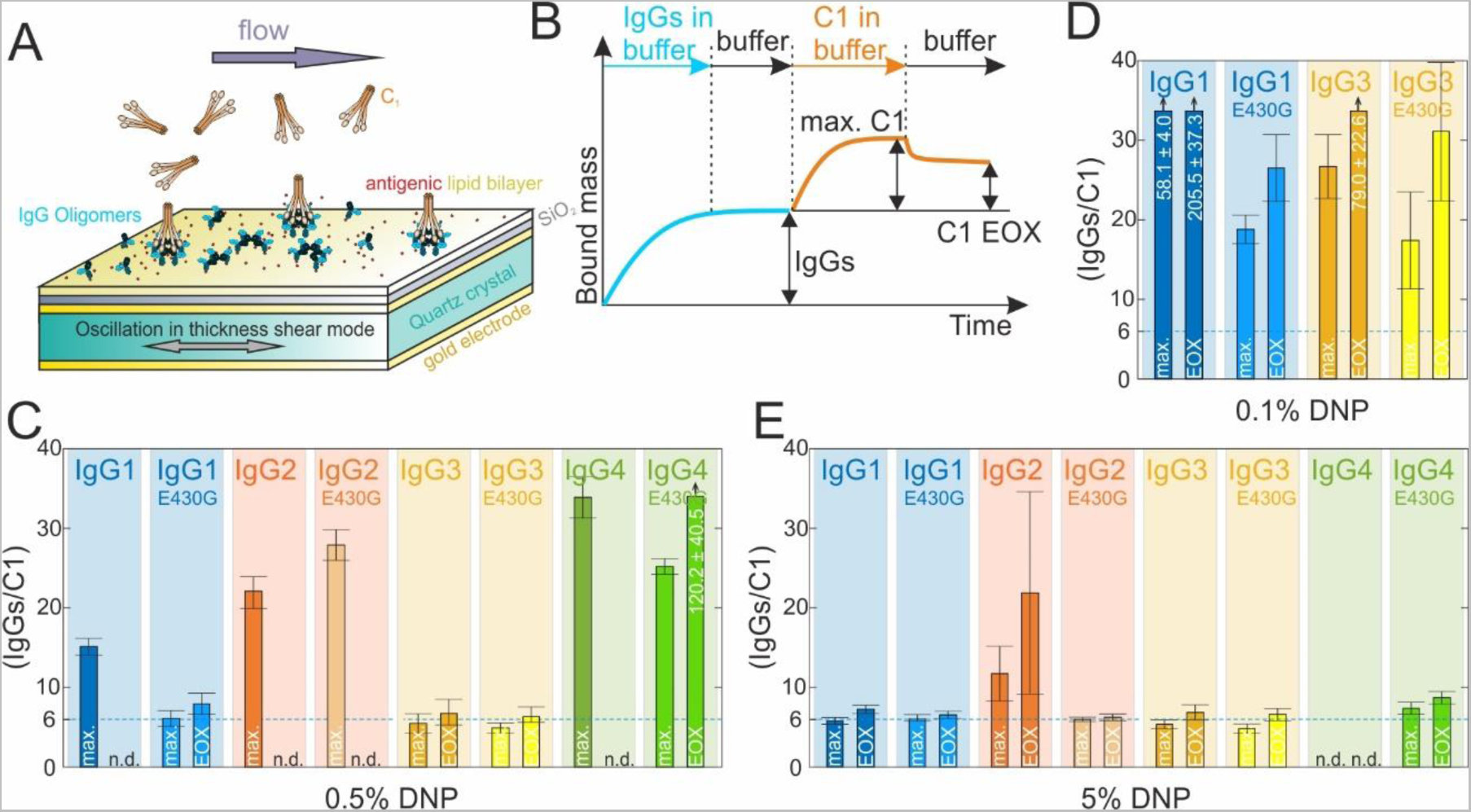
Impact of antigen surface density on C1 recruitment efficiencies of different IgG subclasses and E430G point mutants. **(A)** Illustration of a QCM experiment with C1 molecules in the running buffer over an IgG opsonized DNP-SLB prepared on a SiO2 coated gold electrode of an oscillating quartz crystal. **(B)** Schematic QCM sensorgram of a typical experiment. For calculation of the recruitment efficiencies, the IgG densities were divided by either the max. C1 binding level or the level at the end of the experiment (EOX). **(C)** C1 recruitment efficiencies obtained for a medium antigen density of 0.5 mol % DNP-labelled lipids in the DNP-SLB. **(D)** C1 recruitment efficiencies of IgG1 and IgG3 variants obtained for a low antigen density of 0.1 mol % DNP-labelled lipids in the DNP-SLB. **(E)** C1 recruitment efficiencies of IgG1 and IgG3 variants obtained for a high antigen density of 5 mol % DNP-labelled lipids in the DNP-SLB. Depicted recruitment efficiencies are means ± s.d..

At a medium antigen density (0.5% DNP, Fig. 5C), C1 recruitment was most efficient for IgG3-DNP, followed by IgG1-DNP, IgG2-DNP, and IgG4-DNP. Stable binding with close to no dissociation was only observed for IgG3-DNP (reflected by a comparable IgGs/C1 ratio at max. and EOX). C1 dissociated completely from IgG1-DNP, IgG2-DNP, and IgG4-DNP (n.d.). Importantly, IgG3-DNP exhibited the lowest IgGs/C1 ratios suggesting that on average ∼ six IgG3s were needed to stably bind a single C1 molecule to the antigenic membrane. The E430G point mutation that increased oligomerization of IgG1-DNP-E430G, IgG2-DNP-E430G, and IgG4-DNP-E430G but left IgG3-DNP-E430G oligomerization unaltered (Fig. 4) had a similar effect on C1 binding: It significantly lowered the average number of IgG1-DNP-E430G molecules needed to stably bind a single C1 to ∼ 8 and increased the max. C1 binding of IgG4-DNP-E430G (thus lowering the respective IgGs/C1 ratio), although to a much lesser extent. IgG3-DNP-E430G did not perform significantly differently from the highly efficient IgG3-DNP, suggesting that similarly to oligomerization, C1 binding was not affected by the mutation. To check whether this still holds true for lower antigen densities, we repeated these experiments for IgG1 and IgG3 (and respective E430g variants) using 0.1% DNP SLBs (Fig. 5D). Compared to 0.5% DNP, C1 recruitment and stable binding was vastly reduced for IgG1-DNP, less so for IgG1-DNP-E430G, however, unlike at 0.5% DNP, IgG3-DNP-E430G was significantly more efficient (∼2.5 times) than IgG3-DNP (max. and EOX) in recruiting C1.

Repeating these experiments at higher antigen densities (Fig. 5E) improves C1 recruitment and stable binding for IgG1-DNP, which can no longer be distinguished from IgG1-DNP-E430G, establishes a stable bound C1 population on IgG2-DNP, and enables highly efficient C1 recruitment to IgG2-DNP-E430G. Surprisingly, unlike at lower antigen densities (and accordingly lower IgG4-DNP densities of ∼ 3 x 10^3^ IgGs/µm^2^) where at least transient C1 binding was detectable on IgG4-DNP, high IgG4-DNP densities (∼ 7 x 10^3^ IgGs/µm^2^) resulting from high antigen densities were found to render antigen-bound IgG4-DNP molecules incapable of interacting with C1, presumably because molecular crowding caused by a high density of bivalently bound IgG4-DNP monomers effectively impedes the formation of higher oligomers capable of stable C1 binding. In contrast, IgG4-DNP-E430G was found to exhibit highly efficient C1 recruitment, with levels comparable to the E430G variants of the other three subclasses. Increasing the antigen density did not further improve C1 recruitment efficiency and stability for any IgG variant below ∼ six IgGs/C1 suggesting that the vast majority of antigen bound IgGs are arranged in hexamers at this highest antigen density. Notably, for each IgG variant a certain minimal antigen density is needed to reach this stable level of six IgGs/C1 whereas the E430G point mutation seems to compensate for low antigen densities across subclasses. We have previously (10) noticed that C1 binds stronger to all subclasses than the C1q recognition molecule alone. We hypothesized that C1r_2_s_2_ proteases enhance the stability of the C1q-IgG complexes by fixing the collagen arms, thereby aligning the gC1q domains in a hexagonal platform that favors binding to IgG oligomers. To further substantiate this hypothesis, we have repeated the above QCM experiments with C1q instead of C1 and indeed found that C1q binding is less efficient in all settings except for IgG1-DNP-E430G which was similarly effective in recruiting C1 and C1q on medium and high antigen densities (Fig. S3C-F).

### Complex kinetics of C1/C1q binding to IgG oligomers

Differences in complement activation among IgG subclasses are conventionally rationalized by their respective monovalent affinities for the gC1q headpieces (4). However as shown above, complement binding and CDC clearly correlates with IgG oligomerization on an antigenic surface and can be maximized even for IgG2 and IgG4 by introduction of a mutation that solely enhances IgG oligomerization, not the affinity for gC1q. It is thus likely that the extend of multivalency of the interaction between C1 and IgG oligomers is the main prerequisite for complement activation, while the monovalent affinity of the respective IgG subclass for gC1q plays a rather subordinate role.

To determine affinities and kinetic rate constants of the interactions between gC1q headpieces within C1/C1q and IgG monomers within IgG hexamers, we employed the triple mutants IgG1-DNP-RGY, IgG2-DNP-RGY, IgG3-DNP-RGY, and IgG4-DNP-RGY which were shown to efficiently associate into IgG hexamers in solution (1, 12, 13) to generate an IgG oligomer distribution with a high relative surface density of IgG hexamers. After incubation of the respective IgG-DNP-RGY variant on DNP-SLBs generated on a QCM chip and removal of solution phase IgGs, C1q was added to the running buffer and the binding signal was followed over time (Fig. 6A, exemplified for IgG1-DNP-RGY (I)). Binding quickly reached saturation (II) (Fig. S4) but only minor dissociation, likely from smaller oligomers that form upon antigen binding was observed when C1q was removed from the running buffer (III). No dissociation-rates could be obtained from such curves, because even though gC1q within a multivalently-bound C1q presumably dissociated from IgG Fcs occasionally, immediate rebinding stabilized the hexavalent IgG-RGY - C1q complexes. To nevertheless induce dissociation, we performed competition experiments in which a high affinity competitor (nanobody C1qNb75 (15)) for gC1q was introduced into the QCM flow cell, which strongly binds to transiently unbound gC1q heads, thereby preventing rebinding and eventually leading to complete dissociation of C1q molecules from IgG-RGY hexamers (Fig. 6A, IV). To analyze this data, we developed a corresponding kinetic model of C1/C1q interacting with IgG oligomers (monomers to hexamers) in the absence (Fig. 6A, II and III) and presence (Fig. 6A, IV) of a competitor binding to gC1q heads and thus blocking their interactions with IgG Fcs (Fig. S5). Since the association and dissociation rate constants (k_on,comp_ and k_off,comp_, respectively) of the competitor are known (15), only three additional parameters are required to describe these interactions: The association and dissociation rate constants of the monovalent IgG-Fc - gC1q interactions (k_on,gC1q_ and k_off,gC1q_, respectively) and a parameter that describes the effective concentration (c_eff_) of an unbound gC1q head within an oligomer-bound C1q molecule (Fig. 6B). We performed competitor concentration series for each IgG subclass and fitted the model to the QCM data to extract the three unknown parameters, where k_on,gC1q_ and k_off,gC1q_ were fit locally for each subclass, and c_eff_ globally over all subclasses (Fig. 6C-F and Table 2). The resulting equilibrium dissociation constants (K_D_) were collectively found to be ∼ 10 times lower than previously reported, suggesting that individual IgGs within an oligomer configuration bind C1q with ∼10 times higher affinity as compared to monovalent IgGs in solution (5). The IgG3-gC1q interaction had the lowest K_D_ (4.1 µM), followed by IgG1 (8.0 µM), but different from what was previously found, followed by IgG4 (11.3 µM) and IgG2 (20.0 µM). Repeating this experimental strategy for the interaction between IgG1 and C1 instead of C1q allowed us to quantify the difference between C1 and C1q binding to IgG oligomers. While the association and dissociation rate constants of gC1q heads within C1 and C1q molecules should not be affected by the addition of C1r_2_s_2_ proteases, we have hypothesized that the C1r_2_s_2_ proteases in C1 enhance the stability of C1q-IgG complexes by confining the motional freedom of gC1q heads to a geometry that favors binding to IgG oligomers, which is equivalent to an increase of the effective concentration (c_eff, C1_) in our kinetic model (Fig. 6B). We have thus used the association and dissociation rate constants obtained from the IgG1-RGY – C1q interaction (Fig. 6A) and only adjusted the effective concentration (c_eff, C1_) to fit the IgG1-RGY – C1 competition concentration series (Fig. 6G, Table 2). We found that the presence of C1r_2_s_2_ proteases increases the effective concentration of an unbound gC1q head within an oligomer-bound C1q molecule from 0.4 to 1.5 mM, which corresponds to a reduction of the effective arm length to which the gC1q head is linked within the C1q molecule from 17.7 to 11.7 nm, assuming a spherical volume.

**Figure 6.**
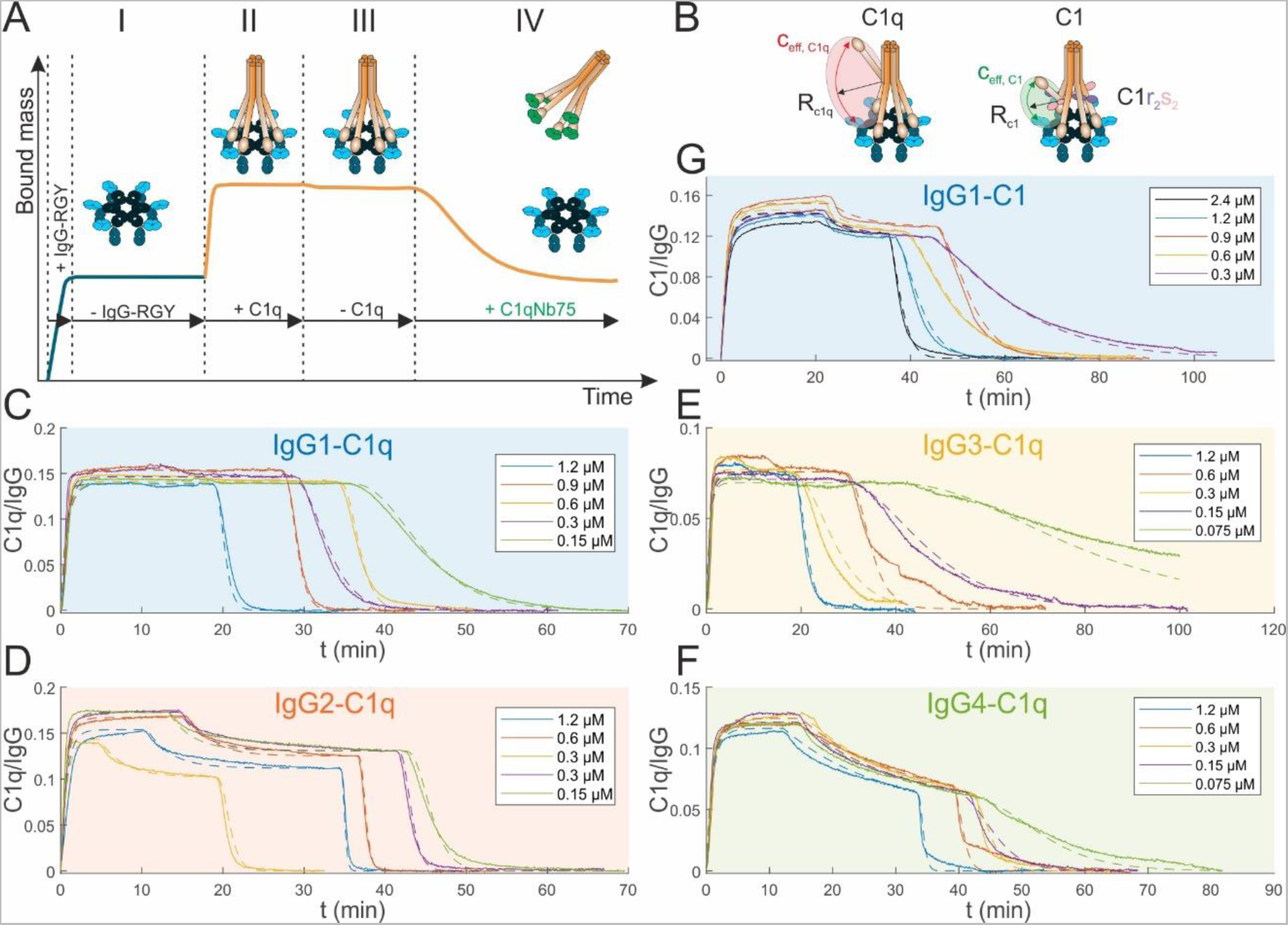
Kinetic analysis of complement C1q and C1 binding to IgG oligomers of different subclasses. **(A)** QCM sensorgram of a typical competition experiment, exemplified by IgG1-DNP-RGY and C1q. After establishing a certain IgG-RGY density on the DNP-SLBs (I) C1q or C1 was added to the QCM running buffer resulting in saturated binding (II; cf. Fig. S4). Removal of C1q/C1 from the running buffer typically induced only minor dissociation (III). Addition of the high-affinity gC1q binding C1qNb75 nanobody induced complete dissociation of C1q/C1 from the underlying IgG oligomers. **(B)** Sketch of C1q/C1 - IgG hexamer complex with an unbound gC1q head. The presence of C1r_2_s_2_ reduces the effective collagen-arm length thereby increasing the local gC1q concentration in C1 as compared to C1q. **(C)** C1qNb75 nanobody concentration series applied to IgG1-DNP-RGY – bound C1q (solid lines). Significant contributions of IgG1-RGY oligomers to C1q binding: 86.2 % in hexamers. **(D)** C1qNb75 nanobody concentration series applied to IgG2-DNP-RGY – bound C1q (solid lines). Significant contributions of IgG2-RGY oligomers to C1q binding: 72.2 % in hexamers, 18.3 % in trimers. **(E)** C1qNb75 nanobody concentration series applied to IgG3-DNP-RGY – bound C1q (solid lines). Significant contributions of IgG3-RGY oligomers to C1q binding: 44.1 % in hexamers. **(F)** C1qNb75 nanobody concentration series applied to IgG4-DNP-RGY – bound C1q (solid lines). Significant contributions of IgG4-RGY oligomers to C1q binding: 30.6 % in hexamers, 22.6 % in trimers. **(G)** C1qNb75 nanobody concentration series applied to IgG1-DNP-RGY – bound C1 (solid lines). Significant contributions of IgG1-RGY oligomers to C1 binding: 73.3 % in hexamers, and 5.7 % in dimers. Dashed lines in (C) – (G) represent fits to our mechanistic model (Fig. S5).

**Table 2.** Parameters obtained from fitting the model (Fig. S3) to the data in Fig. 6B-F. Effective arm lengths were calculated from the effective concentrations by modeling the respective volume as a sphere. Values in brackets are 95% confidence intervals.

|  | IgG1 | IgG2 | IgG3 | IgG4 |
| --- | --- | --- | --- | --- |
| $k_{on, gC1q} (M^{-1}s^{-1})$ | **4.2** (3.2, 5.5) $\times 10^4$ | **5.1** (3.2, 8.3) $\times 10^4$ | **4.5** (2.4, 10.2) $\times 10^4$ | **5.0** (3.3, 9.3) $\times 10^4$ |
| $k_{off, gC1q} (s^{-1})$ | **0.33** (0.22, 0.65) | **1.01** (0.26, 1.74) | **0.18** (0.09, 0.48) | **0.56** (0.45, 1.0) |
| $K_D (\mu M)$ | **7.9** (5.2, 12.1) | **19.8** (4.8, 40.4) | **4.0** (2.6, 6.2) | **11.2** (6.7, 18.1) |
| $C_{eff, C1q} (mM)$ | **0.43** (0.26, 0.81) | | | |
| $C_{eff, C1} (mM)$ | **1.9** (1.7, 2.0) | | | |
| $R_{C1q} (nm)$ | **17.7** (14.3, 20.9) | | | |
| $R_{C1} (nm)$ | **10.8** (10.6, 11.2) | | | |

### IgG oligomerization and resulting C1 binding governs complement dependent lysis of DNP coated vesicles

To link our observations of subclass specific IgG oligomerization, complement C1 recruitment and affinities on DNP-SLBs directly to terminal complement activation, we performed DNP-labeled liposomal vesicle-based complement lysis assays (7) employing the same anti-DNP IgG variants used in our HS-AFM and QCM studies (Fig. 7). Fitting an agonist/response curve to the vesicle lysis data yielded EC50 values (antibody concentration inducing half-maximal lysis) for every IgG variant (Table 3). EC50 was smallest (4.1 nM) for IgG1-DNP, followed by IgG3-DNP (12.5 nM), IgG2-DNP (31.5 nM) and IgG4-DNP with no significant lysis even at the highest IgG4-DNP concentration. The E430G point mutation lowered the EC50 values of IgG1-E430G-DNP (1.6 nM), IgG2-E430G-DNP (12.8 nM), and IgG4-E430G-DNP (4.4 nM) relative to the respective parental IgG, but the EC50 of IgG3-E430G-DNP (12.6 nM) remained essentially unchanged compared to IgG3-DNP (12.5 nM), which is consistent with our QCM experiments performed at the same medium antigen density of 0.5% DNP (Fig. 5C). Triple mutants IgG-RGY-DNP induced strongest complement dependent lysis in all subclassed except for IgG4-RGY which was found less efficient than IgG4-E430G-DNP.

**Figure 7.**
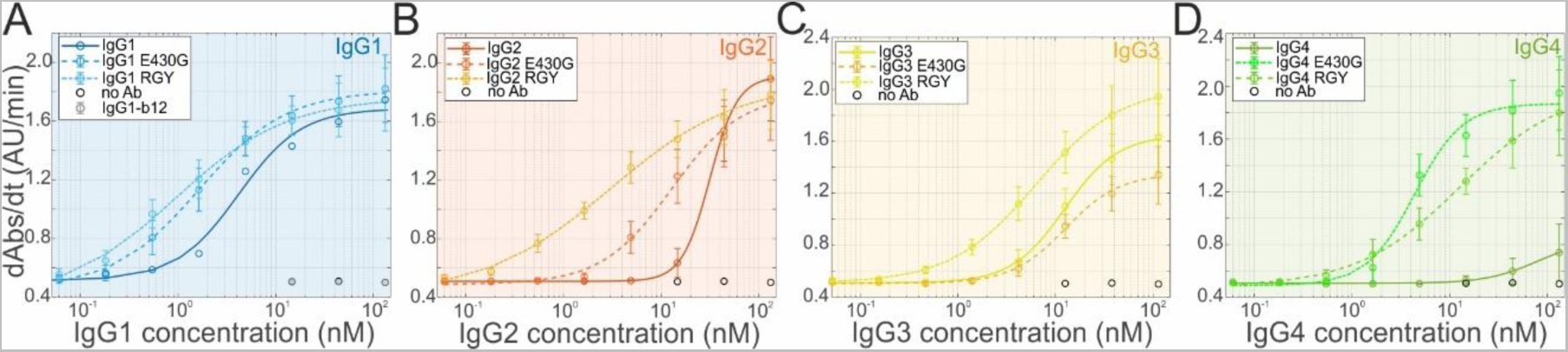
Complement-mediated lysis of DNP-coated liposomes induced by anti-DNP IgG variants. **(A-D)** The ability of graded amounts of IgG1-4 variants to induce activation of complement in human serum is assessed by the magnitude of complement-mediated liposome lysis expressed as the change in absorbance at 340 nm (Abs) per second (dAbs/dt). Each data point represents means ± s.d. from n=3 independent experiments. Lines are fits of dose-response curves to the data, EC50 values are displayed in Table 3.

**Table 3.** EC50 (antibody concentration inducing half-maximal lysis) values for lysis of antibody-opsonized DNP-coated vesicles obtained from fitting a dose-response curve of the type 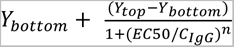 to the CDC data, Fig. 7. Values in brackets are 95% confidence intervals.

|  |  |  |  |
| --- | --- | --- | --- |
|  | IgG1 | IgG1-E430G | IgG1-RGY |
| EC50 (nM) | **4.1** (0.5, 7.8) | **1.6** (1.1, 2.1) | **0.9** (0.4, 1.4) |
|  | IgG2 | IgG2-E430G | IgG2-RGY |
| EC50 (nM) | **31.5** (31.1, 31.8) | **12.8** (7.4, 18.2) | **3.0** (1.5, 4.5) |
|  | IgG3 | IgG3-E430G | IgG3-RGY |
| EC50 (nM) | **12.5** (10.5, 14.4) | **12.6** (9.8, 15.5) | **6.3** (5.7, 6.8) |
|  | IgG4 | IgG4-E430G | IgG4-RGY |
| EC50 (nM) | **n.s.** | **4.4** (2.4, 6.3) | **12.6** (9.7, 15.6) |

## Discussion

The classical pathway of complement activation plays major roles in pathophysiological processes such as infection, inflammation, autoimmunity, or transplant rejection. Its activation depends strongly on the IgG subclass, with IgG1 and IgG3 being able to efficiently trigger activation (16), while IgG2 and IgG4 are usually much less efficient or activate complement only under certain conditions such as IgG2 on high densities of surface antigens like bacterial polysaccharides (10, 17, 18). IgG4 was even reported to form only small immune complexes incapable of complement activation, thereby effectively inhibiting complement activation through competing with IgG1 for targets on cell surfaces (19). The ability to activate complement is associated with the binding affinity of C1q to the respective subclass (5, 16, 20), but also downstream events such as C4b deposition might be differentially affected (9, 16). We have previously shown that antigen bound IgG1 may exist as a distribution of monomers to hexamers. Only tetramers, pentamers, and hexamers could trigger complement-dependent cytotoxicity (CDC), since at least four gC1q headpieces were required to bind to adjacent subunits within these IgG1 oligomers (7). This configuration is consistent with cryo-electron tomography (cryo-ET) images of C1-IgG1 complexes assembled in solution, showing a distribution of complexes with four, five, or six gC1q headpieces adjacently bound to IgG1 hexamers (2), and with cryo-ET images of C1-IgG1 complexes bound to antigenic liposomes, exhibiting the strongest electron densities for four adjacent gC1q headpieces (1). IgM pentamers with one J-chain and IgM hexamers without J-chain adopt very similar structures, where the distances between residues for gC1q binding are identical to the respective distances in IgG1 hexamers (3) representing similar stereochemical danger patterns recognized by C1q. More recently, cryo-ET images of the C1-IgG3 complex revealed similar Fc platforms with all 6 gC1q headpieces bound to IgG3 hexamers (9). These studies suggest that the binding of only two or three adjacent gC1q headpieces to IgG1 or IgG3 oligomers does not result in sufficient compaction of the C1q arms to induce the conformational rearrangements that allow C1r to activate C1s (2). Given these similarities in C1 binding and activation between IgG1, IgG3 and IgM, we here hypothesized that it is primarily an IgG’s ability to oligomerize on antigenic surfaces to form sufficiently large multivalent targets for C1, that ultimately determines if complement activation is initiated or not.

We here demonstrate that indeed all four IgG subclasses are capable of oligomerization, and that the resulting valency of IgG oligomers determines their capacity to multivalently bind C1 which in turn is directly linked to complement activation and CDC. We first evaluated the potency of anti-CD20 and anti-CD52 IgG subclass variants to kill different tumor cells in CDC assays (Fig. 1), by fitting dose-response curves to the CDC data, yielding EC50 values for the respective IgG variants (Table 1). IgG1 and IgG3 were comparably effective in all experimental settings, while IgG2 and IgG4 hardly induced CDC, reproducing the relative CDC efficacies known from literature (16). The E430G point mutation significantly decreased EC50 values for CDC in all cell lines and for all IgG subclasses tested (although less effectively for IgG3), suggesting that it increases IgG2-4 oligomerization in a comparable way as it does for IgG1 (7, 8).

We then quantified IgG oligomerization on antigenic membranes consisting of a supported lipid bilayer model system that contained DNP-labeled lipids in HS-AFM experiments (Fig. 4). Incubation of these DNP-SLBs with equal concentrations of different IgG subclass variants allowed us to collect and compare oligomer distributions for each IgG variant. We found that the abundance of large IgG oligomers was dependent on the IgG subclass - IgG1 was most effective in forming these, followed by IgG3, IgG2 and IgG4. Notably, while IgG4 only formed a minor fraction of pentamers and no hexamers, IgG2 did so much more efficiently, albeit still less than IgG1 and IgG3. The introduction of the E430G point mutation increased the abundance of large IgG oligomers and thus multivalent targets for C1 in all subclasses except in IgG3-E430G-DNP which stayed at the level of IgG3-DNP under these conditions.

Remarkably, IgG3 hexamers extended up to 15 nm above the lipid membrane, further than the other subclasses (10-11 nm), but lower than in a recently proposed structural model based on cryo-ET of IgG3 hexamers protruding up to 19 nm from the membrane. The maximum height of these IgG3 hexamers was found to be associated with a regular array-like arrangement of bivalently bound Fabs that are tightly packed below the Fc platform (9). HS-AFM imaging of IgG3 hexamers revealed additional, moderately mobile lower structures surrounding the central Fc platform which were absent in case of IgG1 hexamers. We interpret these structures as bivalently antigen-bound IgG3 Fab_2_ (Fig. 3) that are connected to the Fc platform via the long flexible hinge of IgG3 but offset from directly below the Fc-platform and thus not arranged in regular arrays as seen in the cryo-ET structures. This is likely caused by the limited antigen and thus antigen-bound Fab_2_ mobility in our model system consisting of gel phase lipids, which effectively hinders Fab_2_ array formation as compared to fluid phase lipid mixture employed in the cryo-ET experiments. Consequently, the Fc-platform is not maximally elevated, but resembles a lower IgG3 hexamer configuration more similar to what was proposed as a transition state to the maximal elevated platform (9). Maximum elevation is thus likely only reached when antigen mobility and antigen lateral sizes allow sufficient proximity of individual Fab_2_ within an IgG3 hexamer to establish intermolecular Fab-Fab contacts responsible for array formation.

C1 binding to antigen-bound IgG oligomers was quantified in QCM experiments (Fig. 5) employing DNP-SLBs containing low, medium, and high antigen densities that resulted in ∼2, 3, and 7 x 10^3^ IgGs/µm^2^ for each IgG subclass variant, respectively. We found that the average number of IgGs bound to surface antigens needed to recruit a single C1 molecule decreased when the epitope density and thus the IgG density was increased, with IgG3 being most effective, followed by IgG1, IgG2 and IgG4. This is in line with recent studies that investigated the influence of antigen densities on CDC, where it was found that IgG3 is superior to IgG1 especially at low antigen densities (18). The recruitment efficiency never went below ∼ 6 IgGs/C1 which hints at the optimal (most stable) complex size being a C1 molecule bound to an IgG hexamer. The E430G variants of each IgG subclass approached this ratio already at lower surface densities than the respective parental IgG, from which we conclude that it was indeed the oligomerization into hexamers (which is enhanced in the E430G variants at a given antigen density) that enabled stable C1 binding rather than the sole density increase of randomly oriented IgGs (which was the same for all IgG variants at a given antigen density). Accordingly, lysis of DNP-labeled vesicles induced by IgG1-E430G-DNP, IgG2-E430G-DNP and IgG4-E430G-DNP improved by the introduction of the mutation, however, similar as observed in the HS-AFM derived IgG oligomer distributions, IgG3-E430G-DNP was not different from IgG3-DNP.

The E430G point mutation increases the flexibility of the IgGs Fc fragments by removing a salt bridge that stabilizes the CH2−CH3 interface packing, normally restraining the domain motility of Fc fragments in the respective IgG (8, 11)). Our current model of IgG oligomerization derived from IgG1 observations (7, 8) suggests that antigen density/availability and affinity on the one hand, and IgG flexibility on the other hand are similarly important for the success of an oligomerization attempt via the so-called vertical pathway, where an antigen-bound IgG (oligomer) recruits an additional IgG from solution via Fc-Fc interactions. In such a situation where the incoming IgG has not yet bound to a surface antigen and is only kept in place by the transient Fc-Fc interaction, the E430G point mutation adds to the overall flexibility of the IgG (mainly given by the hinge flexibility) which is crucial for reaching and binding to nearby antigens. This in turn leads to more successful oligomerization attempts, larger IgG oligomers and thus more efficient C1 binding at lower antigen densities compared to the respective parental IgGs that lack that additional flexibility. This is in line with CDC data from B cells of chronic lymphocytic leukemia (CLL) patients showing a 10-fold range of different CD20 expression levels. Anti-CD20 IgG1-7D8-E430G was similarly effective throughout the whole CD20 range, while CDC mediated by IgG1-7D8 decreased significantly at lower CD20 levels (21). The effect of E430G on already highly flexible IgG3 was thus only apparent at lowest antigen densities where IgG3-E430G exhibited slightly improved C1 recruitment with respect to IgG3, which is likely the reason why we did not observe a similar difference in our HS-AFM characterization of IgG3 oligomer distributions (Fig. 4) and liposomal vesicle-based complement lysis assays (Fig. 7), which were both done at higher antigen densities. IgG4 was found to bind C1 only transiently at medium antigen densities which was not sufficient to induce lysis of DNP-labeled vesicles (Fig. 7) and did not recruit any C1 at high antigen and thus high IgG4 surface densities (Fig. 5), suggesting while at medium antigen densities oligomerization into smaller oligomers occurred, molecular crowding at elevated IgG4 densities effectively hindered oligomer formation so that no binding sites for C1 were generated at all, supporting the notion of IgG4 acting as a “blocking antibody” (19).

While IgG2 did not induce CDC of tumor cells, large IgG2 oligomers (Fig. 4), C1 binding (Fig. 5) and complement mediated lysis of DNP-labeled vesicles (Fig. 7), although less potent than IgG1 and IgG3, have been observed. We attribute this to the 2-10 fold higher antigen surface densities in our model system as compared to the cell surface CD20 densities on DAUDI and RAJI cells used in CDC assays (8.1 x 10^3^ DNP molecules/ µm^2^ resulting from 0.5 mol % DNP-cap-DPPE content (22) compared to 1.4 – 3.6 x 10^3^ CD20 molecules/µm^2^ on DAUDI cells and 0.7 – 1.7 x 10^3^ CD20 molecules/µm^2^ on RAJI and WIEN cells; based on a diameter of 5-8 µm (12) and respective CD20 expression levels per cell (11)), highlighting the restriction of complement activation by IgG2 to targets expressing elevated densities of surface antigens like bacterial polysaccharides (10, 17).

The relative capacity of the four IgG subclasses to activate complement is often argued based on their respective monovalent affinities for the gC1q headpieces (4), as characterized in analytical ultracentrifugation experiments (5). However, we found that complement binding and CDC clearly correlate with the ability of an IgG variant to oligomerize on an antigenic surface and can be maximized by introduction of a mutation that enhances oligomerization but leaves the actual binding epitopes for gC1q unaltered (23). This holds true even for IgG2 and IgG4 that have, according to (5), the lowest affinity for gC1q. To clarify the role of affinity for gC1q vs. oligomerization ability of the different IgG subclasses, we developed a mechanistic model of the interactions between C1/C1q and IgG monomers to hexamers and used this model to analyze QCM sensorgrams of the interaction between IgG hexamers and C1/C1q in the presence of a high affinity competitor for gC1q (Fig. S5). This allowed us to not only extract K_D_ values of the monovalent interactions responsible for the strong multivalent binding of C1/C1q to IgG hexamers, but also association and dissociation rate constants that govern the kinetics of the interactions. To adequately describe the multivalent binding between C1/C1q and IgG oligomers, it was necessary to introduce a parameter that accounts for the effective concentration (c_eff_) of an unbound gC1q head within an oligomer-bound C1 or C1q molecule, which was found to be four times larger in C1 than in C1q, quantifying for the first time the contribution of C1r_2_s_2_ proteases to the stability of the IgG hexamer C1 complex (10). The effective arm length of the collagen-like C1q segments resulting from these concentrations (Table 2) agree reasonably well with the respective arm length measured from either the N-terminal stalk region to the gC1q head (C1q) or from the position of the C1r_2_s_2_ heterotrimer to the gC1q head in C1 (2). The cryo-ET structures of soluble C1-IgG1 hexamer complexes reported in this latter study were categorized as separate classes with four, five, or six gC1q domains in contact with Fc platforms. The class with six gC1q heads bound appeared most frequently, followed by four and five gC1q domains bound, suggesting that C1 bound with an even number of gC1q heads are more stable (per bond) than the ones bound with an odd number. In contrast, our model does not differentiate between even and odd numbers of bound gC1q heads and thus predicts a monotonic increase of the number of gC1q heads bound to the Fc platform within an ensemble of C1-IgG1 hexamer complexes. Introduction of an additional parameter accounting for a pair-wise cooperativity in gC1q binding (practically by introducing two different c_eff_ that apply alternating) would in principle enable such a non-monotonic behavior as observed in cryo-ET structures, however, given that our model already reasonably fits our QCM data, and only marginal improvements could be expected from a more complex model, we retained the simpler model.

Remarkably, the resulting K_D_ values of the monovalent interactions between all IgGs and gC1q were approximately 10 times lower than previously reported, and the resulting ranking was IgG3 > IgG1 > IgG4 > IgG2, in order of decreasing affinity for gC1q. Previous attempts to determine these monovalent affinities for all subclasses were performed in solution (5) where the conformation of the IgGs is considerably different (24) from ordered IgG oligomers observed on antigenic membranes (1–3). Accordingly, affinities determined in this way can differ largely from affinities determined when one of the interaction partners is confined in a certain conformation such as IgGs within antigen-bound IgG oligomers. Other experimental designs that involve the unspecific adsorption of IgGs to solid surfaces and thus their confinement into 2D (25), resulted in estimates for K_D_ values in the same order of magnitude as determined from our QCM experiments. Remarkably, although intact IgG4 did not interact with C1 in theses assays, its Fc fragment was able to interact comparable to IgG1 Fc fragments, with K_D_ values for gC1q of ∼ 4–7 µM (estimated from the presented dose response curves). These are in excellent agreement with the K_D_ values determined here for the interaction of IgG1/IgG4 within hexamers and gC1q (8.0 and 11.3 µM, respectively) suggesting a certain similarity between the Fc platform of IgG hexamers and individual Fc fragments. While the binding site for gC1q located in the Fc-CH2 domain close to the hinge is likely largely masked by the Fabs within IgGs that float freely in solution (especially when the hinge is short and relatively stiff as in case of IgG4), the fixation of at least one of the Fabs within an IgG hexamer through antigen binding and the orientation of the Fc fragment within the Fc platform will likely increase the accessibility of the gC1q binding site, similarly as the enzymatic removal of the Fabs does when Fc fragments are generated from IgGs.

By extending the expression for the functional affinity (also termed avidity) of two monovalent interactions that are linked via a flexible linker molecule (8, 26), we can estimate the functional affinity between C1(q) and IgG subclass oligomers of different sizes n = 1 – 6 via

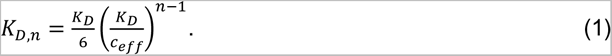

Putting the subclass specific gC1q – IgG K_D_ values and the effective concentration for C1 or C1q (Table S2 and Fig. 6B) into this equation allows us to derive functional affinity constants for all 4 IgG subclasses and all oligomer sizes, as displayed in Fig. 8A and B for C1 and C1q, respectively. The thermodynamic derivation (Eq. 1) thereby reproduces the K_D_ values obtained from directly fitting a Langmuir isotherm to the simulation of the concentration dependent equilibrium binding of C1/C1q to IgG oligomers (Fig. 8C, exemplified for C1 binding to IgG1 monomers - hexamers) based on our mechanistic model Fig. S5. Provided that the requirements for C1 activation by IgG1, i.e. that at least four gC1q heads must bind to adjacent IgGs within an oligomer (7) apply to all four subclasses, the functional affinities of the resulting complexes (i.e. tetramers and larger) are in the pM range and below, so that at physiological C1 concentrations of ∼170 nM, strong binding and activation should occur as soon as oligomers of at least the size of tetramers are present on a target cell surface. It is thus the ability to oligomerize into large enough IgG oligomers on a certain target cell that decides if a certain antibody subclass/mutant variant is capable of activating C1. Consequently, the actual monovalent affinity for gC1q heads likely plays a rather subordinate role, except in governing the transient unbinding/rebinding rates of gC1q within the complexes, which could affect C1r/C1s activation differently in the individual subclasses. Removal of C1r_2_s_2_ from IgG-bound C1 by serum concentrations of human C1-ersterase inhibitor (C1-INH) was reported to induce vast dissociation of C1q from IgG2, while C1q bound to IgG1 was much less affected, and C1q remained bound to IgG3 at essentially the same level as complete C1 (10). In our model, removal of C1r_2_s_2_ corresponds to a switch from c_eff_ = 1.5 mM (C1) to c_eff_ = 0.4 mM (C1q) leading to a loss in functional affinity that indeed results in a strong dissociation of C1q from IgG2 trimers and tetramers as compared to C1, whereas IgG1 and IgG3 oligomers are less or not at all affected, Fig. S6.

**Figure 8.**
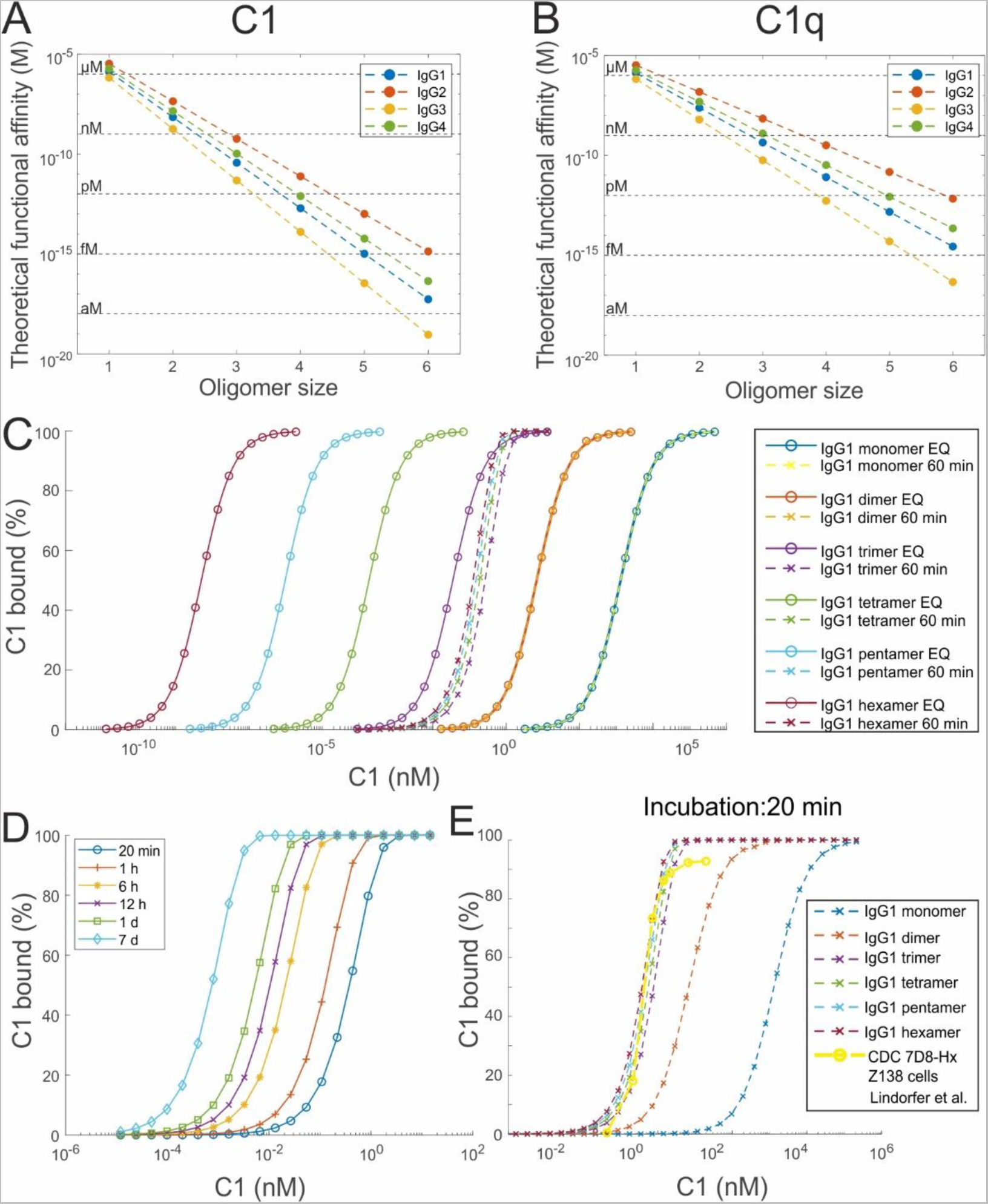
C1/C1q binding to differently sized IgG subclass oligomers. **(A-B)** Comparison of functional affinities of C1 (A) and C1q (B) to differently sized IgG subclass oligomers determined from kinetic simulations (until equilibrium is reached, dots) and from the thermodynamic approach (dashed lines), using the parameters from Table 2, respectively. **(C)** Concentration dependency of C1 binding to IgG1 monomers to hexamers at equilibrium (circles) and after an incubation time of 60 min (crosses). **(D)** Influence of incubation time on C1 binding to IgG1 hexamers. **(E)** Concentration dependency of C1 binding to IgG1 monomers to hexamers after an incubation time of 20 min. Comparison to CDC data of anti-CD20 antibody IgG1-7D8-E430G bound to Z138 cells taken from (29).

Unlike the sub-pM affinity of the C1-activating complexes for C1 determined here, several authors report apparent K_D_ values for C1 binding to antibody-opsonized cells in the 1-100 nM range (1, 27). For instance, analysis of C1q binding signals obtained from real-time cell-binding assays detected two distinct binding modes of C1q characterized by a strong and a weak interaction component with K_D_ values of 0.07-2.3 and 5-70 nM respectively (28). These values coincide with the K_D_ values determined here (Fig. 8B) for the interaction of C1q with IgG1 trimers (0.44 nM) and IgG1 dimers (23.9 nM), suggesting that in this case the majority of C1q is recruited via these smaller IgG oligomers, which are, however, unable to activate complement (7). Yet, C1 binding levels needed to kill substantially amounts of antibody opsonized cells have been reported to be well below saturating C1 levels. For instance, C3b deposition and CDC are already maximal for anti-CD20 antibody ofatumumab (OFA) -opsonized RAJI cells at less than 10 % of its maximum C1 binding level (27), and the oligomerization enhanced E430G point mutant of the anti-CD20 IgG1-7D8 was found to induce maximum CDC already at 1 - 2 nM C1, while much higher C1 concentrations were needed to fully saturate C1 binding on the used B-cell lines (29).

This is in line with our finding that IgGs exist in the form of oligomer populations on antigenic surfaces with higher oligomers being much less abundant than smaller ones (Fig. 4), and that the functional affinity increases distinctly with oligomer size. IgG1 oligomers capable of activating C1, bind C1 with a sub-pM functional affinity (tetramers and larger, Fig. 8A), and, provided that the incubation time of IgG opsonized cells with C1 allowed the reaction to reach equilibrium for a given C1 concentration (Fig. 8D exemplified for an IgG hexamer), binding to these oligomers saturates and induces maximum CDC well below maximum C1 binding levels that are dominated by the binding to the much higher abundant IgG dimers and trimers in the used experimental C1 concentration range. Consequently, the maximum C1 binding level is not a good predictor for complement activation since the maximum is dominated by contributions from non-C1 activating, small IgG oligomers. Contributions of C1 binding to IgG1 monomers (K_D_ = 1.3 µM) are not observed in such binding curves, since the C1 concentration is usually not increased beyond the physiological concentration of ∼ 170 nM. Importantly, typical incubation times (60 min in (1), 20 min in (29)) in combination with low C1 concentrations in the pM regime do not allow the system to reach equilibrium and thus do not lead to saturating binding of C1 to the complement activating higher IgG1 oligomers (Fig. 8C and D). In this case, the respective C1 binding levels that are reached at the end of the incubation time generate a C1 concentration-dependency for all the large C1-activating IgG oligomers which is similar to what was observed in CDC lysis experiments performed at such low C1 concentrations and short incubation times (29). The CDC vs. C1 concentration curve of the most potent anti-CD20 IgG1 used in this study perfectly overlaps with this theoretical prediction of C1 binding to IgG1 tetramers to hexamers (Fig. 8E). Lower densities of large IgG1 oligomers in combination with the presence of differing amounts of complement regulatory proteins such as CD46, CD55, and CD59 likely further shift the concentration dependency to higher C1 concentrations, which might explain the differences observed among different antibody variants and cell lines (29), however this will be subject to future investigations.

Additionally to the saturating binding of C1 to complement-activating IgG tetramers to hexamers, *in vivo* C1 concentrations (0.17 µM in human serum) will also lead to saturating C1 binding to antigen-bound IgG dimers and trimers, but not to IgG monomers, which might be relevant for non-complement activation related functions of C1 such as its role as ligand for cell surface receptors on innate and adaptive immune cells (30), or as competitors for Fcγ receptors that exhibit K_D_ values for IgGs in a similar range (31). Taken together, we present comprehensive insights into the complex interplay of innate antibody properties and multivalent IgG-C1q/C1 interactions, that determine IgG subclass specific differences in complement recruitment and activation. Furthermore, we suggest how complement C1 binding to large oligomers is related to the outcome of CDC, which might offer a way to characterize oligomer distributions and the impact of complement regulatory proteins on different target cells. These insights will allow IgG subclass-specific differences to be specifically adapted and exploited in future immunotherapies.

## Materials and Methods

### DNP labeled liposomes

DNP-labeled liposomes consisting of 1,2-dipalmitoyl-sn-glycero-3-phosphocholine (DPPC), 1,2-dipalmitoyl-sn-glycero-3-phosphoethanolamine (DPPE) and 1,2-dipalmitoyl-sn-glycero-3-phosphoethanolamine-N-[6-[(2,4-dinitrophenyl)amino]hexanoyl] (DNP-cap-DPPE) were used to generate supported lipid bilayers (SLBs) on mica and SiO2 substrates. The lipids were purchased from Avanti Polar Lipids, mixed at different ratios of DPPC:DPPE:DNP-cap-DPPE (90:5:5, 90:9.5:0.5, and 90:9.9:0.1 molar ratios), and dissolved in a 2:1 mixture of chloroform and methanol. After the solvents were rotary-evaporated for 30 min, the lipids were again dissolved in chloroform, which was then rotary-evaporated for 30 min. Drying was completed at a high vacuum pump for 2 h. The lipids were dissolved in 500 μL Milli-Q H2O while immersed in a water bath at 60 °C, flooded with argon, and sonicated for 3 min at 60 °C to create small unilamellar vesicles. These were diluted to 2 mg/mL in buffer #1 (10 mM HEPES, 150 mM NaCl, 2 mM CaCl2, pH 7.4) and frozen for storage using liquid N2.

### High-Speed Atomic Force Microscopy (HS-AFM)

HS-AFM (32) (SS-NEX, RIBM Ltd., Ibaraki, JP) was conducted in tapping mode at RT (room temperature, 25°C) in liquid, with typical free amplitudes of 1.5-2.5 nm and amplitude setpoints larger than 90%. Silicon nitride cantilevers with electron-beam deposited tips USC-F1.2-k0.15 (NanoWorld® AG, Neuchatel, CH), nominal spring constants of 0.15 N m^−1^, resonance frequencies around 500 kHz, and a quality factor of approx. 2 in liquids were used.

DNP labeled supported lipid bilayers (DNP-SLBs) for HS-AFM were prepared on muscovite mica. The liposomes were incubated on the freshly cleaved surface (0.5 mg/ml in buffer #1), placed in a humidity chamber to prevent evaporation, and heated to 60°C for 30 min. Then the temperature was gradually cooled down to RT within 30 min, followed by exchanging the solution with buffer #1. After 10 min of equilibration at RT, and 15 more buffer exchanges, the SLB was ready for imaging. To passivate any exposed mica, SLBs were incubated with 330 nM IgG1-b12 (isotype control antibody against HIV-1 gp120) (33) for 10 min before the molecules of interest were added. The height distributions (Figures 2) were obtained after incubating DNP-SLBs with 33.3 nM of the respective IgG variant for 5 min. The sample was then imaged with a frame size of 400 x 400 nm² in buffer #1, and for each distinct location on the sample (n > 15) a short video was recorded (2 - 5 frames). The position of the individual particles and complexes within these videos were tracked using the ImageJ (NIH) plugin Mosaic Suite Particle Tracker (34) and correlated with the respective height information via a MATLAB (The MathWorks Inc., MA, US) script developed in-house. The height average was determined for each particle over the course of the video, pooled for each IgG variant, and plotted as probability density functions (pdfs) and histograms. The oligomer distributions on DNP-SLBs were analyzed in a two-step process: Individual particle dimensions were determined by HS-AFM, and their oligomeric state was further confirmed via their decay pattern determined in subsequent forced dissociation experiments (7). In brief, molecules were scanned in a non-disrupting manner to gauge their number, height, and shape. Subsequently, the scanning force exerted by the HS-AFM cantilever tip is increased (by decreasing the setpoint-amplitude) to dissociate oligomers into their constituent IgGs. Geometric parameters and oligomer decay patterns are combined to assign each IgG assembly its oligomeric state.

### Quartz Crystal Microbalance (QCM)

QCM experiments were done using a two-channel QCM-I system (MicroVacuum). AT cut SiO2-coated quartz crystals with a diameter of 14.0 mm and a resonance frequency of 5 MHz were used (Quartz Pro AB). All sensorgrams were recorded on the first, third, and fifth harmonic frequencies. The data shown are related to the third harmonic. Before each set of experiments, the SiO_2_-coated crystals were cleaned by immersion in 2% sodium dodecyl sulfate (SDS) for 30 min, followed by thorough rinsing with Milli-Q H2O. The chips were dried in a gentle stream of N_2_ and oxidized using air plasma (4 min at 80 W; Diener electronic GmbH & Co. KG, Ebhausen, DE), then mounted in the measurement chamber. The sensor surface and the fluid system was further cleaned by a flow of 2% SDS at 250 μL min^−1^ for 5 min, followed by Milli-Q H_2_O at 250 μL min^−1^ for 5 min directly before the measurements. Before lipid incubation, the flow was stopped and the measurement chamber was heated up to 45 °C, left on this temperature for a few minutes and cooled down again to 22 °C for equilibration. To generate DNP-SLBs on QCM chips, the DPPC:DPPE:DNP-cap-DPPE liposome stock solution was heated to 60 °C for 30 min and then slowly cooled to RT within 30 min. The solution was ready for injection after dilution to 200 µg/mL with buffer #1. DNP-SLB formation was typically complete after 30 min at μL min^−1^, after which the flow medium was changed to buffer #1 and a second heat cycle was started, followed by equilibration in buffer #1. Finally, a control injection with 15.2 nM C1q (Complement Technology Inc., TX, US), was performed to check for imperfections in the DNP-SLBs. Competition experiments (Fig. 6) were performed in either buffer #1 (C1q) or buffer #2 (155 mM NaCl, 9.6 mM HEPES, 1.9 mM CaCl_2_, 1.8 mM sodium acetate, 1.8 mM EACA, 0.4 mM benzamidin HCl, 0.4 mM EDTA, 1.4% glycerol, pH 7.4; C1).

Raw-sensorgrams (in Hz *vs.* time) were converted to molecule densities on the QCM chip (molecules/µm^2^ *vs.* time) by determining the bound mass according to the Sauerbrey equation (35) which relates the change in resonance frequency Δf of a quartz crystal oscillating in thickness shear mode to the mass adsorbed on its surface Δm by the relation 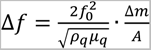. Here f_0_ is the fundamental resonance frequency, and ρ_q_ and μ_q_ are the crystal density (2.648 g cm^−3^) and its shear modulus (2.947 10^11^ g cm^−1^ s^2^), respectively. The resulting masses were corrected for the effect of trapped water according to Höök *et al.*(36). Determination of molecule densities further requires the effective surface area of the DNP-SLBs, which may exceed the actual working surface area of the QCM chip due to residual, only partially fused (and thus non-flat) vesicles. We corrected the active surface area accordingly by comparing the theoretical mass of a flat DNP-SLB covering the working surface (average molecular weight of 729.3 Da for DPPC:DPPE=9:1; head-group area of 0.626 nm²,(37) 0.5 nm buffer layer between SiO_2_ and DNP-SLB) to the actual observed mass, and modeling the excess as buffer filled half spheres with a diameter ∼ 35 nm (determined by HS-AFM imaging).

### Data fitting and simulations

Fitting of dose-response curves to CDC and vesicle-based complement lysis assays, fitting of QCM senorgrams, and simulation of C1/C1q binding to IgG oligomers according to our mechanistic model (Fig. S5, Supporting Information) was done in MATLAB (The MathWorks Inc., MA, US). For the latter, the ODE15s and fminsearch nonlinear solver were used. Confidence intervals of fit parameters were determined according to (38, 39).

### Cell lines

FreeStyle™ Expi293F™ cells were cultured in FreeStyle™ 293 expression medium according to the manufacturer’s instructions (Invitrogen). Additional cell lines were obtained from the American Type Culture Collection (ATCC). Raji and Daudi (human CD20-positive Burkitt’s lymphoma) cells were cultured in RPMI 1640 medium (Lonza), supplemented with 10% (v/v) heat-inactivated Donor Bovine Serum with Iron (DBSI; Life Technologies). Wien-133 (human CD52-positive Burkitt’s lymphoma) cells were cultured in Iscove’s Modified Dulbeco’s Medium (IMDM) with HEPES and L-Glutamine (Lonza), supplemented with 10% (v/v) heat-inactivated DBSI. All cell lines were maintained at 37°C in a 5% (v/v) CO_2_ humidified incubator.

### Construction, expression, and purification of antibody variants

Antibody heavy-chain (HC) expression vectors were constructed by inserting de novo synthesized (Geneart) codon optimized HC coding regions into expression vector pcDNA3.3 (Invitrogen). The HC coding regions consisted of the VH regions of mAbs 7D8 (human CD20-specific [Teeling et al., 2004]), Campath (human CD52-specific [Hale et al., 1983], G2a2 (DNP-specific [White et al., 1996]) or b12 (HIV-1 gp120-specific [Roben et al., 1994]), genetically fused to the CH regions of wild-type human IgG1*03, IgG2*01, IgG3*01 or IgG4*01 or one of the mutants variants containing the E430G point mutation [de Jong et al., 2016] or the RGY (E345R-E430G-S440Y) triple-mutation [Diebolder et al., 2014] (EU numbering conventions are used throughout the manuscript). Likewise, separate light-chain expression vectors were constructed by inserting the corresponding VL coding regions in frame with the CL coding regions of the human (J00241) kappa light chain into expression vector pcDNA3.3.

All antibodies were produced under serum-free conditions by co-transfecting relevant heavy and light chain expression vectors in FreeStyle™ Expi293F™ cells, using ExpiFectamine™ 293 (LifeTechnologies), according to the manufacturer’s instructions. IgG1, IgG2 and IgG4 antibody variants were purified by protein A affinity chromatography (MabSelect SuRe; GE Health Care), dialyzed overnight to PBS and filter-sterilized over 0.2-µM dead-end filters. Alternatively, IgG3 antibody variants were purified by protein G affinity chromatography (GE Health Care). Purity was determined by CE-SDS and concentration was measured by absorbance at 280 nm (specific extinction coefficients were calculated for each protein). Batches of purified antibody were tested by high-performance size-exclusion chromatography (HP-SEC) for aggregates or degradation products and shown to be at least 95% monomeric. Purified antibodies were stored at 2-8°C.

### Complement dependent cytotoxicity (CDC) assay

The capacity of anti-CD20 and anti-CD52 antibodies to induce CDC was assessed by pre-incubating Raji (1×10^5^ cells), Daudi (1×10^5^ cells) or Wien133 target cells (3×10^5^ cells) in assay buffer (RPMI medium containing 0.1% (w/v) BSA) at 21°C for 15 min with serial diluted antibodies. Pooled human serum (20% (v/v)) was added as a source of complement and cells were incubated at 37°C for an additional 45 min. Cells were then put on ice and viability was determined by staining with propidium iodide (PI) and detected using an iQue screener (Intellicyt). Percentage lysis was calculated using the following formula: % lysis = (experimental release (fluorescence) – spontaneous release without antibody (fluorescence))/ (maximal release of IgG1 (fluorescence) – spontaneous release without antibody (fluorescence)) x 100.

### DNP-labeled liposomal vesicle-based complement lysis assay

For the DNP-labeled liposomal vesicle-based complement lysis assay we used the Wako Auto Kit (Wako Pure Chemical, Chuo-Ku Osaka, Japan) with an adapted protocol to be able to test the anti-DNP variants used. In a regular 96 ELISA flat bottom plate 15% normal human serum (Sanquin, Amsterdam, The Netherlands) was added to the liposome mixture (R1 from the kit) containing G6PDH and a serial dilution of antibody (20 x concentrated) and incubated for 5 minutes at room temperature. Then 33.5% custom substrate mix (containing 24 mM D-glucose-6-phospate solution, 9 mM beta-Nicotinamide adenine dinucleotide, 20.6 mM lactose monohydrate, 9.8 mM NaOH (Sigma), 1:1 mixed with maleate buffer (R2a from the kit)) was added and incubated at room temperature and measured kinetically on an Envision microplate reader (PerkinElmer, Waltham, MA) at 340 nm for 20 min. The change in absorbance (Abs per second was calculated from the linear part of the curve) dAbs/dt was plotted against log transformed antibody concentration.

## Acknowledgments

We thank J.I. Meesters, K Beslmuller and C. Le for technical assistance. J.P. acknowledges support the Austrian Science Fund (FWF, Grant No. P33958 and P34164) and the Federal State of Upper Austria as a part of the FH Upper Austria Center of Excellence for Technological Innovation in Medicine (TIMed Center).

## Author Contributions

J.S., A.F.L, F.J.B and J.P. designed research. N.F., J.S., E.G.F.B. performed experiments and analyzed data. J.P. supervised the project, developed the model, did data analysis, prepared the figures and wrote the paper. All authors read and commented on the paper.

