## Supporting Information for "Complement activation by IgG subclasses is governed by their ability to oligomerize upon antigen binding"

### Supporting Information for
Complement activation by IgG subclasses is governed by their ability to oligomerize upon antigen binding

Nikolaus Frischauf[1], Jürgen Strasser[1], Ellen G.F. Borg[2], Aran F. Labrijn[2], Frank J. Beurskens[2], Johannes Preiner[1]

[1] University of Applied Sciences Upper Austria, Linz, Austria

[2] Genmab, Utrecht, Netherlands

*Johannes Preiner

**This PDF file includes:**

Supporting text
Figures S1 to S6
Legends for Movies S1 to S4
SI References

**Other supporting materials for this manuscript include the following:**

Movies S1 to S4

#### Supporting Information Text

##### Mechanistic model of C1/C1q binding to IgG oligomers

We have deduced the system of rate equations governing the time course of C1q/C1 binding/dissociation to/from different IgG oligomers from the kinetic scheme given in Fig. S5. The model is based on the assumption that after the removal of IgG from solution, the resulting IgG oligomer distributions do not change over time, which is justified by the observation that after dissociation of C1q/C1 induced by a competitor, the subsequent C1q/C1 incubations on the very same IgG-opsonized DNP-SLBs lead to similarly shaped sensorgrams and reached similar binding levels as the respective preceding experiments. For the different states of IgG-oligomer – C1q/C1 – competitor complexes we introduced a three-digit index notation $y_{nmo}$, where *n, m, and o* depict the size of the IgG oligomer, the number of available (i.e. not yet bound to an IgG or a competitor) gC1q heads, and the number of competitor-bound gC1q heads within a C1q/C1 molecule, respectively. For example, $y_{512}$ (Fig. S5B, 3rd row, 3rd column) represents a C1 molecule bound to an IgG pentamer (5), having one gC1q head still unbound and thus available (1), and two gC1q heads occupied by a competitor (2), and consequently 6-1-2 = 3 gC1q heads already bound to the IgG pentamer. The corresponding rate equation that governs the time evolution of this state can be obtained by adding up all flux from neighboring states into/from $y_{512}$. Starting from left to right and from bottom to top we get:

$$\frac{d\boldsymbol{y}_{512}}{dt} = \left(2 \cdot \boldsymbol{k}_{on,comp}\boldsymbol{C}_C\boldsymbol{y}_{521} - 2 \cdot \boldsymbol{k}_{off,comp}\boldsymbol{y}_{512}\right) - \left(1 \cdot \boldsymbol{k}_{on,comp}\boldsymbol{C}_C\boldsymbol{y}_{512} - 3 \cdot \boldsymbol{k}_{off,comp}\boldsymbol{y}_{503}\right) - \left(2 \cdot \frac{1}{3} \cdot \boldsymbol{k}_{on}\boldsymbol{c}_{eff}\boldsymbol{y}_{512} - 4 \cdot \boldsymbol{k}_{off}\boldsymbol{y}_{502}\right) + \left(3 \cdot \frac{2}{4} \cdot \boldsymbol{k}_{on}\boldsymbol{c}_{eff}\boldsymbol{y}_{522} - 3 \cdot \boldsymbol{k}_{off}\boldsymbol{y}_{512}\right)$$

The numerical pre-factors of the respective expressions account for the respective probabilities that a certain transition may occur, by assuming that only adjacent gC1q heads may bind to adjacent IgGs within an oligomer. IgG oligomers without C1/C1q bound are depicted by single-digit indexes, e.g. $y_5$ for an IgG pentamer.

The complete set of rate equations then reads (next page):

#### I. C1/C1q binding to IgG hexamers (Fig. S5A):

**1st row (one gC1q head bound):**

$$\frac{d\boldsymbol{y}_{650}}{dt} = -\left(5\cdot \boldsymbol{k}_{on,comp}\boldsymbol{C}_C\boldsymbol{y}_{650} - 1\cdot \boldsymbol{k}_{off,comp}\boldsymbol{y}_{641}\right) - \left(5\cdot \boldsymbol{k}_{on}\boldsymbol{c}_{eff}\boldsymbol{y}_{650} - 2\cdot \boldsymbol{k}_{off}\boldsymbol{y}_{640}\right) + \left(36\cdot \boldsymbol{k}_{on}\boldsymbol{C}_{C1}\boldsymbol{y}_{6} - \boldsymbol{k}_{off}\boldsymbol{y}_{650}\right)$$

$$\frac{d\boldsymbol{y}_{641}}{dt} = \left(5\cdot \boldsymbol{k}_{on,comp}\boldsymbol{C}_C\boldsymbol{y}_{650} - 1\cdot \boldsymbol{k}_{off,comp}\boldsymbol{y}_{641}\right) - \left(4\cdot \boldsymbol{k}_{on,comp}\boldsymbol{C}_C\boldsymbol{y}_{641} - 2\cdot \boldsymbol{k}_{off,comp}\boldsymbol{y}_{632}\right) - \left(4\cdot \boldsymbol{k}_{on}\boldsymbol{c}_{eff}\boldsymbol{y}_{641} - 2\cdot \boldsymbol{k}_{off}\boldsymbol{y}_{631}\right) + \left( -\boldsymbol{k}_{off}\boldsymbol{y}_{641}\right)$$

$$\frac{d\boldsymbol{y}_{632}}{dt} = \left(4\cdot \boldsymbol{k}_{on,comp}\boldsymbol{C}_C\boldsymbol{y}_{641} - 2\cdot \boldsymbol{k}_{off,comp}\boldsymbol{y}_{632}\right) - \left(3\cdot \boldsymbol{k}_{on,comp}\boldsymbol{C}_C\boldsymbol{y}_{632} - 3\cdot \boldsymbol{k}_{off,comp}\boldsymbol{y}_{623}\right) - \left(3\cdot \boldsymbol{k}_{on}\boldsymbol{c}_{eff}\boldsymbol{y}_{632} - 2\cdot \boldsymbol{k}_{off}\boldsymbol{y}_{622}\right) + \left( -\boldsymbol{k}_{off}\boldsymbol{y}_{632}\right)$$

$$\frac{d\boldsymbol{y}_{623}}{dt} = \left(3\cdot \boldsymbol{k}_{on,comp}\boldsymbol{C}_C\boldsymbol{y}_{632} - 3\cdot \boldsymbol{k}_{off,comp}\boldsymbol{y}_{623}\right) - \left(2\cdot \boldsymbol{k}_{on,comp}\boldsymbol{C}_C\boldsymbol{y}_{623} - 4\cdot \boldsymbol{k}_{off,comp}\boldsymbol{y}_{614}\right) - \left(2\cdot \boldsymbol{k}_{on}\boldsymbol{c}_{eff}\boldsymbol{y}_{623} - 2\cdot \boldsymbol{k}_{off}\boldsymbol{y}_{613}\right) + \left( -\boldsymbol{k}_{off}\boldsymbol{y}_{623}\right)$$

$$\frac{d\boldsymbol{y}_{614}}{dt} = \left(2\cdot \boldsymbol{k}_{on,comp}\boldsymbol{C}_C\boldsymbol{y}_{623} - 4\cdot \boldsymbol{k}_{off,comp}\boldsymbol{y}_{614}\right) - \left(1\cdot \boldsymbol{k}_{on,comp}\boldsymbol{C}_C\boldsymbol{y}_{614} - 5\cdot \boldsymbol{k}_{off,comp}\boldsymbol{y}_{605}\right) - \left(1\cdot \boldsymbol{k}_{on}\boldsymbol{c}_{eff}\boldsymbol{y}_{614} - 2\cdot \boldsymbol{k}_{off}\boldsymbol{y}_{604}\right) + \left( -\boldsymbol{k}_{off}\boldsymbol{y}_{614}\right)$$

$$\frac{d\boldsymbol{y}_{605}}{dt} = \left(1\cdot \boldsymbol{k}_{on,comp}\boldsymbol{C}_C\boldsymbol{y}_{614} - 5\cdot \boldsymbol{k}_{off,comp}\boldsymbol{y}_{605}\right) + \left( -\boldsymbol{k}_{off}\boldsymbol{y}_{605}\right)$$

**2nd row (two gC1q heads bound):**

$$\frac{d\boldsymbol{y}_{640}}{dt} = -\left(4\cdot \boldsymbol{k}_{on,comp}\boldsymbol{C}_C\boldsymbol{y}_{640} - 1\cdot \boldsymbol{k}_{off,comp}\boldsymbol{y}_{631}\right) - \left(4\cdot \boldsymbol{k}_{on}\boldsymbol{c}_{eff}\boldsymbol{y}_{640} - 3\cdot \boldsymbol{k}_{off}\boldsymbol{y}_{630}\right) + \left(5\cdot \boldsymbol{k}_{on}\boldsymbol{c}_{eff}\boldsymbol{y}_{650} - 2\cdot \boldsymbol{k}_{off}\boldsymbol{y}_{640}\right)$$

$$\frac{d\boldsymbol{y}_{631}}{dt} = \left(4\cdot \boldsymbol{k}_{on,comp}\boldsymbol{C}_C\boldsymbol{y}_{640} - 1\cdot \boldsymbol{k}_{off,comp}\boldsymbol{y}_{631}\right) - \left(3\cdot \boldsymbol{k}_{on,comp}\boldsymbol{C}_C\boldsymbol{y}_{631} - 2\cdot \boldsymbol{k}_{off,comp}\boldsymbol{y}_{622}\right) - \left(3\cdot \boldsymbol{k}_{on}\boldsymbol{c}_{eff}\boldsymbol{y}_{631} - 3\cdot \boldsymbol{k}_{off}\boldsymbol{y}_{621}\right) + \left(4\cdot \boldsymbol{k}_{on}\boldsymbol{c}_{eff}\boldsymbol{y}_{641} - 2\cdot \boldsymbol{k}_{off}\boldsymbol{y}_{631}\right)$$

$$\frac{d\boldsymbol{y}_{622}}{dt} = \left(3\cdot \boldsymbol{k}_{on,comp}\boldsymbol{C}_C\boldsymbol{y}_{631} - 2\cdot \boldsymbol{k}_{off,comp}\boldsymbol{y}_{622}\right) - \left(2\cdot \boldsymbol{k}_{on,comp}\boldsymbol{C}_C\boldsymbol{y}_{622} - 3\cdot \boldsymbol{k}_{off,comp}\boldsymbol{y}_{613}\right) - \left(2\cdot \boldsymbol{k}_{on}\boldsymbol{c}_{eff}\boldsymbol{y}_{622} - 3\cdot \boldsymbol{k}_{off}\boldsymbol{y}_{612}\right) + \left(3\cdot \boldsymbol{k}_{on}\boldsymbol{c}_{eff}\boldsymbol{y}_{632} - 2\cdot \boldsymbol{k}_{off}\boldsymbol{y}_{622}\right)$$

$$\frac{d\boldsymbol{y}_{613}}{dt} = \left(2\cdot \boldsymbol{k}_{on,comp}\boldsymbol{C}_C\boldsymbol{y}_{622} - 3\cdot \boldsymbol{k}_{off,comp}\boldsymbol{y}_{613}\right) - \left(1\cdot \boldsymbol{k}_{on,comp}\boldsymbol{C}_C\boldsymbol{y}_{613} - 4\cdot \boldsymbol{k}_{off,comp}\boldsymbol{y}_{604}\right) - \left(1\cdot \boldsymbol{k}_{on}\boldsymbol{c}_{eff}\boldsymbol{y}_{613} - 3\cdot \boldsymbol{k}_{off}\boldsymbol{y}_{603}\right) + \left(2\cdot \boldsymbol{k}_{on}\boldsymbol{c}_{eff}\boldsymbol{y}_{623} - 2\cdot \boldsymbol{k}_{off}\boldsymbol{y}_{613}\right)$$

$$\frac{d\boldsymbol{y}_{604}}{dt} = \left(1\cdot \boldsymbol{k}_{on,comp}\boldsymbol{C}_C\boldsymbol{y}_{613} - 4\cdot \boldsymbol{k}_{off,comp}\boldsymbol{y}_{604}\right) + \left(1\cdot \boldsymbol{k}_{on}\boldsymbol{c}_{eff}\boldsymbol{y}_{614} - 2\cdot \boldsymbol{k}_{off}\boldsymbol{y}_{604}\right)$$

**3rd row (three gC1q heads bound):**

$$\frac{d\boldsymbol{y}_{630}}{dt} = -\left(3\cdot \boldsymbol{k}_{on,comp}\boldsymbol{C}_C\boldsymbol{y}_{630} - 1\cdot \boldsymbol{k}_{off,comp}\boldsymbol{y}_{621}\right) - \left(3\cdot \boldsymbol{k}_{on}\boldsymbol{c}_{eff}\boldsymbol{y}_{630} - 4\cdot \boldsymbol{k}_{off}\boldsymbol{y}_{620}\right) + \left(4\cdot \boldsymbol{k}_{on}\boldsymbol{c}_{eff}\boldsymbol{y}_{640} - 3\cdot \boldsymbol{k}_{off}\boldsymbol{y}_{630}\right)$$

$$\frac{d\boldsymbol{y}_{621}}{dt} = \left(3\cdot \boldsymbol{k}_{on,comp}\boldsymbol{C}_C\boldsymbol{y}_{630} - 1\cdot \boldsymbol{k}_{off,comp}\boldsymbol{y}_{621}\right) - \left(2\cdot \boldsymbol{k}_{on,comp}\boldsymbol{C}_C\boldsymbol{y}_{621} - 2\cdot \boldsymbol{k}_{off,comp}\boldsymbol{y}_{612}\right) - \left(2\cdot \boldsymbol{k}_{on}\boldsymbol{c}_{eff}\boldsymbol{y}_{621} - 4\cdot \boldsymbol{k}_{off}\boldsymbol{y}_{611}\right) + \left(3\cdot \boldsymbol{k}_{on}\boldsymbol{c}_{eff}\boldsymbol{y}_{631} - 3\cdot \boldsymbol{k}_{off}\boldsymbol{y}_{621}\right)$$

$$\frac{d\boldsymbol{y}_{612}}{dt} = \left(2\cdot \boldsymbol{k}_{on,comp}\boldsymbol{C}_C\boldsymbol{y}_{621} - 2\cdot \boldsymbol{k}_{off,comp}\boldsymbol{y}_{612}\right) - \left(1\cdot \boldsymbol{k}_{on,comp}\boldsymbol{C}_C\boldsymbol{y}_{612} - 3\cdot \boldsymbol{k}_{off,comp}\boldsymbol{y}_{603}\right) - \left(1\cdot \boldsymbol{k}_{on}\boldsymbol{c}_{eff}\boldsymbol{y}_{612} - 4\cdot \boldsymbol{k}_{off}\boldsymbol{y}_{602}\right) + \left(2\cdot \boldsymbol{k}_{on}\boldsymbol{c}_{eff}\boldsymbol{y}_{622} - 3\cdot \boldsymbol{k}_{off}\boldsymbol{y}_{612}\right)$$

$$\frac{d\boldsymbol{y}_{603}}{dt} = \left(1\cdot \boldsymbol{k}_{on,comp}\boldsymbol{C}_C\boldsymbol{y}_{612} - 3\cdot \boldsymbol{k}_{off,comp}\boldsymbol{y}_{603}\right) + \left(1\cdot \boldsymbol{k}_{on}\boldsymbol{c}_{eff}\boldsymbol{y}_{613} - 3\cdot \boldsymbol{k}_{off}\boldsymbol{y}_{603}\right)$$

**4th row (four gC1q heads bound):**

$$\frac{dy_{620}}{dt} = -\left(2 \cdot k_{on,comp} C_C y_{620} - 1 \cdot k_{off,comp} y_{611}\right) - \left(2 \cdot k_{on} c_{eff} y_{620} - 5 \cdot k_{off} y_{610}\right) + \left(3 \cdot k_{on} c_{eff} y_{630} - 4 \cdot k_{off} y_{620}\right)$$

$$\frac{dy_{611}}{dt} = \left(2 \cdot k_{on,comp} C_C y_{620} - 1 \cdot k_{off,comp} y_{611}\right) - \left(1 \cdot k_{on,comp} C_C y_{611} - 2 \cdot k_{off,comp} y_{602}\right) - \left(1 \cdot k_{on} c_{eff} y_{611} - 5 \cdot k_{off} y_{601}\right) + \left(2 \cdot k_{on} c_{eff} y_{621} - 4 \cdot k_{off} y_{611}\right)$$

$$\frac{dy_{602}}{dt} = \left(1 \cdot k_{on,comp} C_C y_{611} - 2 \cdot k_{off,comp} y_{602}\right) + \left(1 \cdot k_{on} c_{eff} y_{612} - 4 \cdot k_{off} y_{602}\right)$$

**5th row (five gC1q heads bound):**

$$\frac{dy_{610}}{dt} = -\left(1 \cdot k_{on,comp} C_C y_{610} - 1 \cdot k_{off,comp} y_{601}\right) - \left(1 \cdot k_{on} c_{eff} y_{610} - 6 \cdot k_{off} y_{600}\right) + \left(2 \cdot k_{on} c_{eff} y_{620} - 5 \cdot k_{off} y_{610}\right)$$

$$\frac{dy_{601}}{dt} = \left(1 \cdot k_{on,comp} C_C y_{610} - 1 \cdot k_{off,comp} y_{601}\right) + \left(1 \cdot k_{on} c_{eff} y_{611} - 5 \cdot k_{off} y_{601}\right)$$

**6th row (six gC1q heads bound):**

$$\frac{dy_{600}}{dt} = +\left(1 \cdot k_{on} c_{eff} y_{610} - 6 \cdot k_{off} y_{600}\right)$$

**Hexamers without C1/C1q:**

$$\frac{dy_6}{dt} = -36 \cdot k_{on} C_{C1} y_6 + k_{off}(y_{650} + y_{641} + y_{632} + y_{623} + y_{614} + y_{605})$$

#### II. C1/C1q binding to IgG pentamers (Fig. S5B):

**1st row (one gC1q head bound):**

$$\frac{dy_{550}}{dt} = -\left(5 \cdot k_{on,comp} C_C y_{550} - 1 \cdot k_{off,comp} y_{541}\right) - \left(4 \cdot \frac{5}{5} \cdot k_{on} c_{eff} y_{550} - 2 \cdot k_{off} y_{540}\right) + (30 \cdot k_{on} C_{C1} y_5 - k_{off} y_{550})$$

$$\frac{dy_{541}}{dt} = \left(5 \cdot k_{on,comp} C_C y_{550} - 1 \cdot k_{off,comp} y_{541}\right) - \left(4 \cdot k_{on,comp} C_C y_{541} - 2 \cdot k_{off,comp} y_{532}\right) - \left(4 \cdot \frac{4}{5} \cdot k_{on} c_{eff} y_{541} - 2 \cdot k_{off} y_{531}\right) + ( - k_{off} y_{541})$$

$$\frac{dy_{532}}{dt} = \left(4 \cdot k_{on,comp} C_C y_{541} - 2 \cdot k_{off,comp} y_{532}\right) - \left(3 \cdot k_{on,comp} C_C y_{532} - 3 \cdot k_{off,comp} y_{523}\right) - \left(4 \cdot \frac{3}{5} \cdot k_{on} c_{eff} y_{532} - 2 \cdot k_{off} y_{522}\right) + ( - k_{off} y_{532})$$

$$\frac{dy_{523}}{dt} = \left(3 \cdot k_{on,comp} C_C y_{532} - 3 \cdot k_{off,comp} y_{523}\right) - \left(2 \cdot k_{on,comp} C_C y_{523} - 4 \cdot k_{off,comp} y_{514}\right) - \left(4 \cdot \frac{2}{5} \cdot k_{on} c_{eff} y_{523} - 2 \cdot k_{off} y_{513}\right) + ( - k_{off} y_{523})$$

$$\frac{dy_{514}}{dt} = \left(2 \cdot k_{on,comp} C_C y_{523} - 4 \cdot k_{off,comp} y_{514}\right) - \left(1 \cdot k_{on,comp} C_C y_{514} - 5 \cdot k_{off,comp} y_{505}\right) - \left(4 \cdot \frac{1}{5} \cdot k_{on} c_{eff} y_{514} - 2 \cdot k_{off} y_{504}\right) + ( - k_{off} y_{514})$$

$$\frac{dy_{505}}{dt} = \left(1 \cdot k_{on,comp} C_C y_{514} - 5 \cdot k_{off,comp} y_{505}\right) + ( - k_{off} y_{505})$$

**2nd row (two gC1q heads bound):**

$$\frac{dy_{540}}{dt} = -\left(4 \cdot k_{on,comp} C_C y_{540} - 1 \cdot k_{off,comp} y_{531}\right) - \left(3 \cdot \frac{4}{4} \cdot k_{on} c_{eff} y_{540} - 3 \cdot k_{off} y_{530}\right) + \left(4 \cdot \frac{5}{5} \cdot k_{on} c_{eff} y_{550} - 2 \cdot k_{off} y_{540}\right)$$

$$\frac{d\boldsymbol{y}_{531}}{dt} = (4\cdot \boldsymbol{k}_{on,comp}\boldsymbol{C}_C\boldsymbol{y}_{540} - 1\cdot \boldsymbol{k}_{off,comp}\boldsymbol{y}_{531}) - (3\cdot \boldsymbol{k}_{on,comp}\boldsymbol{C}_C\boldsymbol{y}_{531} - 2\cdot \boldsymbol{k}_{off,comp}\boldsymbol{y}_{522}) - \left(3\cdot\frac{3}{4}\cdot \boldsymbol{k}_{on}\boldsymbol{c}_{eff}\boldsymbol{y}_{531} - 3\cdot \boldsymbol{k}_{off}\boldsymbol{y}_{521}\right) + \left(4\cdot\frac{4}{5}\cdot \boldsymbol{k}_{on}\boldsymbol{c}_{eff}\boldsymbol{y}_{541} - 2\cdot \boldsymbol{k}_{off}\boldsymbol{y}_{531}\right)$$

$$\frac{d\boldsymbol{y}_{522}}{dt} = (3\cdot \boldsymbol{k}_{on,comp}\boldsymbol{C}_C\boldsymbol{y}_{531} - 2\cdot \boldsymbol{k}_{off,comp}\boldsymbol{y}_{522}) - (2\cdot \boldsymbol{k}_{on,comp}\boldsymbol{C}_C\boldsymbol{y}_{522} - 3\cdot \boldsymbol{k}_{off,comp}\boldsymbol{y}_{513}) - \left(3\cdot\frac{2}{4}\cdot \boldsymbol{k}_{on}\boldsymbol{c}_{eff}\boldsymbol{y}_{522} - 3\cdot \boldsymbol{k}_{off}\boldsymbol{y}_{512}\right) + \left(4\cdot\frac{3}{5}\cdot \boldsymbol{k}_{on}\boldsymbol{c}_{eff}\boldsymbol{y}_{532} - 2\cdot \boldsymbol{k}_{off}\boldsymbol{y}_{522}\right)$$

$$\frac{d\boldsymbol{y}_{513}}{dt} = (2\cdot \boldsymbol{k}_{on,comp}\boldsymbol{C}_C\boldsymbol{y}_{522} - 3\cdot \boldsymbol{k}_{off,comp}\boldsymbol{y}_{513}) - (1\cdot \boldsymbol{k}_{on,comp}\boldsymbol{C}_C\boldsymbol{y}_{513} - 4\cdot \boldsymbol{k}_{off,comp}\boldsymbol{y}_{504}) - \left(3\cdot\frac{1}{4}\cdot \boldsymbol{k}_{on}\boldsymbol{c}_{eff}\boldsymbol{y}_{513} - 3\cdot \boldsymbol{k}_{off}\boldsymbol{y}_{503}\right) + \left(4\cdot\frac{2}{5}\cdot \boldsymbol{k}_{on}\boldsymbol{c}_{eff}\boldsymbol{y}_{523} - 2\cdot \boldsymbol{k}_{off}\boldsymbol{y}_{513}\right)$$

$$\frac{d\boldsymbol{y}_{504}}{dt} = (1\cdot \boldsymbol{k}_{on,comp}\boldsymbol{C}_C\boldsymbol{y}_{513} - 4\cdot \boldsymbol{k}_{off,comp}\boldsymbol{y}_{504}) + \left(4\cdot\frac{1}{5}\cdot \boldsymbol{k}_{on}\boldsymbol{c}_{eff}\boldsymbol{y}_{514} - 2\cdot \boldsymbol{k}_{off}\boldsymbol{y}_{504}\right)$$

**3rd row (three gC1q heads bound):**

$$\frac{d\boldsymbol{y}_{530}}{dt} = -(3\cdot \boldsymbol{k}_{on,comp}\boldsymbol{C}_C\boldsymbol{y}_{530} - 1\cdot \boldsymbol{k}_{off,comp}\boldsymbol{y}_{521}) - \left(2\cdot\frac{3}{3}\cdot \boldsymbol{k}_{on}\boldsymbol{c}_{eff}\boldsymbol{y}_{530} - 4\cdot \boldsymbol{k}_{off}\boldsymbol{y}_{520}\right) + \left(3\cdot\frac{4}{4}\cdot \boldsymbol{k}_{on}\boldsymbol{c}_{eff}\boldsymbol{y}_{540} - 3\cdot \boldsymbol{k}_{off}\boldsymbol{y}_{530}\right)$$

$$\frac{d\boldsymbol{y}_{521}}{dt} = (3\cdot \boldsymbol{k}_{on,comp}\boldsymbol{C}_C\boldsymbol{y}_{530} - 1\cdot \boldsymbol{k}_{off,comp}\boldsymbol{y}_{521}) - (2\cdot \boldsymbol{k}_{on,comp}\boldsymbol{C}_C\boldsymbol{y}_{521} - 2\cdot \boldsymbol{k}_{off,comp}\boldsymbol{y}_{512}) - \left(2\cdot\frac{2}{3}\cdot \boldsymbol{k}_{on}\boldsymbol{c}_{eff}\boldsymbol{y}_{521} - 4\cdot \boldsymbol{k}_{off}\boldsymbol{y}_{511}\right) + \left(3\cdot\frac{3}{4}\cdot \boldsymbol{k}_{on}\boldsymbol{c}_{eff}\boldsymbol{y}_{531} - 3\cdot \boldsymbol{k}_{off}\boldsymbol{y}_{521}\right)$$

$$\frac{d\boldsymbol{y}_{512}}{dt} = (2\cdot \boldsymbol{k}_{on,comp}\boldsymbol{C}_C\boldsymbol{y}_{521} - 2\cdot \boldsymbol{k}_{off,comp}\boldsymbol{y}_{512}) - (1\cdot \boldsymbol{k}_{on,comp}\boldsymbol{C}_C\boldsymbol{y}_{512} - 3\cdot \boldsymbol{k}_{off,comp}\boldsymbol{y}_{503}) - \left(2\cdot\frac{1}{3}\cdot \boldsymbol{k}_{on}\boldsymbol{c}_{eff}\boldsymbol{y}_{512} - 4\cdot \boldsymbol{k}_{off}\boldsymbol{y}_{502}\right) + \left(3\cdot\frac{2}{4}\cdot \boldsymbol{k}_{on}\boldsymbol{c}_{eff}\boldsymbol{y}_{522} - 3\cdot \boldsymbol{k}_{off}\boldsymbol{y}_{512}\right)$$

$$\frac{d\boldsymbol{y}_{503}}{dt} = (1\cdot \boldsymbol{k}_{on,comp}\boldsymbol{C}_C\boldsymbol{y}_{512} - 3\cdot \boldsymbol{k}_{off,comp}\boldsymbol{y}_{503}) + \left(3\cdot\frac{1}{4}\cdot \boldsymbol{k}_{on}\boldsymbol{c}_{eff}\boldsymbol{y}_{513} - 3\cdot \boldsymbol{k}_{off}\boldsymbol{y}_{503}\right)$$

**4th row (four gC1q heads bound):**

$$\frac{d\boldsymbol{y}_{520}}{dt} = -(2\cdot \boldsymbol{k}_{on,comp}\boldsymbol{C}_C\boldsymbol{y}_{520} - 1\cdot \boldsymbol{k}_{off,comp}\boldsymbol{y}_{511}) - \left(1\cdot\frac{2}{2}\cdot \boldsymbol{k}_{on}\boldsymbol{c}_{eff}\boldsymbol{y}_{520} - 5\cdot \boldsymbol{k}_{off}\boldsymbol{y}_{510}\right) + \left(2\cdot\frac{3}{3}\cdot \boldsymbol{k}_{on}\boldsymbol{c}_{eff}\boldsymbol{y}_{530} - 4\cdot \boldsymbol{k}_{off}\boldsymbol{y}_{520}\right)$$

$$\frac{d\boldsymbol{y}_{511}}{dt} = (2\cdot \boldsymbol{k}_{on,comp}\boldsymbol{C}_C\boldsymbol{y}_{520} - 1\cdot \boldsymbol{k}_{off,comp}\boldsymbol{y}_{511}) - (1\cdot \boldsymbol{k}_{on,comp}\boldsymbol{C}_C\boldsymbol{y}_{511} - 2\cdot \boldsymbol{k}_{off,comp}\boldsymbol{y}_{502}) - \left(1\cdot\frac{1}{2}\cdot \boldsymbol{k}_{on}\boldsymbol{c}_{eff}\boldsymbol{y}_{511} - 5\cdot \boldsymbol{k}_{off}\boldsymbol{y}_{501}\right) + \left(2\cdot\frac{2}{3}\cdot \boldsymbol{k}_{on}\boldsymbol{c}_{eff}\boldsymbol{y}_{521} - 4\cdot \boldsymbol{k}_{off}\boldsymbol{y}_{511}\right)$$

$$\frac{d\boldsymbol{y}_{502}}{dt} = (1\cdot \boldsymbol{k}_{on,comp}\boldsymbol{C}_C\boldsymbol{y}_{511} - 2\cdot \boldsymbol{k}_{off,comp}\boldsymbol{y}_{502}) + \left(2\cdot\frac{1}{3}\cdot \boldsymbol{k}_{on}\boldsymbol{c}_{eff}\boldsymbol{y}_{512} - 4\cdot \boldsymbol{k}_{off}\boldsymbol{y}_{502}\right)$$

**5th row (five gC1q heads bound):**

$$\frac{d\boldsymbol{y}_{510}}{dt} = -(1\cdot \boldsymbol{k}_{on,comp}\boldsymbol{C}_C\boldsymbol{y}_{510} - 1\cdot \boldsymbol{k}_{off,comp}\boldsymbol{y}_{501}) + \left(1\cdot\frac{2}{2}\cdot \boldsymbol{k}_{on}\boldsymbol{c}_{eff}\boldsymbol{y}_{520} - 5\cdot \boldsymbol{k}_{off}\boldsymbol{y}_{510}\right)$$

$$\frac{d\boldsymbol{y}_{501}}{dt} = (1\cdot \boldsymbol{k}_{on,comp}\boldsymbol{C}_C\boldsymbol{y}_{510} - 1\cdot \boldsymbol{k}_{off,comp}\boldsymbol{y}_{501}) + \left(1\cdot\frac{1}{2}\cdot \boldsymbol{k}_{on}\boldsymbol{c}_{eff}\boldsymbol{y}_{511} - 5\cdot \boldsymbol{k}_{off}\boldsymbol{y}_{501}\right)$$

**Pentamers without C1/C1q:**

$$\frac{d\boldsymbol{y}_5}{dt} = -30\cdot \boldsymbol{k}_{on}\boldsymbol{C}_{C1}\boldsymbol{y}_5 + \boldsymbol{k}_{off}(\boldsymbol{y}_{550} + \boldsymbol{y}_{541} + \boldsymbol{y}_{532} + \boldsymbol{y}_{523} + \boldsymbol{y}_{514} + \boldsymbol{y}_{505})$$

#### III. C1/C1q binding to IgG Tetramers (Fig. S5C):

**1st row (one gC1q head bound):**

$$\frac{d\boldsymbol{y}_{450}}{dt} = -\left(5\cdot\boldsymbol{k}_{on,comp}\boldsymbol{C}_{C}\boldsymbol{y}_{450} - 1\cdot\boldsymbol{k}_{off,comp}\boldsymbol{y}_{441}\right) - \left(3\cdot\frac{5}{5}\cdot\boldsymbol{k}_{on}\boldsymbol{c}_{eff}\boldsymbol{y}_{450} - 2\cdot\boldsymbol{k}_{off}\boldsymbol{y}_{440}\right) + \left(24\cdot\boldsymbol{k}_{on}\boldsymbol{C}_{C1}\boldsymbol{y}_{4} - \boldsymbol{k}_{off}\boldsymbol{y}_{450}\right)$$

$$\frac{d\boldsymbol{y}_{441}}{dt} = \left(5\cdot\boldsymbol{k}_{on,comp}\boldsymbol{C}_{C}\boldsymbol{y}_{450} - 1\cdot\boldsymbol{k}_{off,comp}\boldsymbol{y}_{441}\right) - \left(4\cdot\boldsymbol{k}_{on,comp}\boldsymbol{C}_{C}\boldsymbol{y}_{441} - 2\cdot\boldsymbol{k}_{off,comp}\boldsymbol{y}_{432}\right) - \left(3\cdot\frac{4}{5}\cdot\boldsymbol{k}_{on}\boldsymbol{c}_{eff}\boldsymbol{y}_{441} - 2\cdot\boldsymbol{k}_{off}\boldsymbol{y}_{431}\right) + \left(-\boldsymbol{k}_{off}\boldsymbol{y}_{441}\right)$$

$$\frac{d\boldsymbol{y}_{432}}{dt} = \left(4\cdot\boldsymbol{k}_{on,comp}\boldsymbol{C}_{C}\boldsymbol{y}_{441} - 2\cdot\boldsymbol{k}_{off,comp}\boldsymbol{y}_{432}\right) - \left(3\cdot\boldsymbol{k}_{on,comp}\boldsymbol{C}_{C}\boldsymbol{y}_{432} - 3\cdot\boldsymbol{k}_{off,comp}\boldsymbol{y}_{423}\right) - \left(3\cdot\frac{3}{5}\cdot\boldsymbol{k}_{on}\boldsymbol{c}_{eff}\boldsymbol{y}_{432} - 2\cdot\boldsymbol{k}_{off}\boldsymbol{y}_{422}\right) + \left(-\boldsymbol{k}_{off}\boldsymbol{y}_{432}\right)$$

$$\frac{d\boldsymbol{y}_{423}}{dt} = \left(3\cdot\boldsymbol{k}_{on,comp}\boldsymbol{C}_{C}\boldsymbol{y}_{432} - 3\cdot\boldsymbol{k}_{off,comp}\boldsymbol{y}_{423}\right) - \left(2\cdot\boldsymbol{k}_{on,comp}\boldsymbol{C}_{C}\boldsymbol{y}_{423} - 4\cdot\boldsymbol{k}_{off,comp}\boldsymbol{y}_{414}\right) - \left(3\cdot\frac{2}{5}\cdot\boldsymbol{k}_{on}\boldsymbol{c}_{eff}\boldsymbol{y}_{423} - 2\cdot\boldsymbol{k}_{off}\boldsymbol{y}_{413}\right) + \left(-\boldsymbol{k}_{off}\boldsymbol{y}_{423}\right)$$

$$\frac{d\boldsymbol{y}_{414}}{dt} = \left(2\cdot\boldsymbol{k}_{on,comp}\boldsymbol{C}_{C}\boldsymbol{y}_{423} - 4\cdot\boldsymbol{k}_{off,comp}\boldsymbol{y}_{414}\right) - \left(1\cdot\boldsymbol{k}_{on,comp}\boldsymbol{C}_{C}\boldsymbol{y}_{414} - 5\cdot\boldsymbol{k}_{off,comp}\boldsymbol{y}_{405}\right) - \left(3\cdot\frac{1}{5}\cdot\boldsymbol{k}_{on}\boldsymbol{c}_{eff}\boldsymbol{y}_{414} - 2\cdot\boldsymbol{k}_{off}\boldsymbol{y}_{404}\right) + \left(-\boldsymbol{k}_{off}\boldsymbol{y}_{414}\right)$$

$$\frac{d\boldsymbol{y}_{405}}{dt} = \left(1\cdot\boldsymbol{k}_{on,comp}\boldsymbol{C}_{C}\boldsymbol{y}_{414} - 5\cdot\boldsymbol{k}_{off,comp}\boldsymbol{y}_{405}\right) + \left(-\boldsymbol{k}_{off}\boldsymbol{y}_{405}\right)$$

**2nd row (two gC1q heads bound):**

$$\frac{d\boldsymbol{y}_{440}}{dt} = -\left(4\cdot\boldsymbol{k}_{on,comp}\boldsymbol{C}_{C}\boldsymbol{y}_{440} - 1\cdot\boldsymbol{k}_{off,comp}\boldsymbol{y}_{431}\right) - \left(2\cdot\frac{4}{4}\cdot\boldsymbol{k}_{on}\boldsymbol{c}_{eff}\boldsymbol{y}_{440} - 3\cdot\boldsymbol{k}_{off}\boldsymbol{y}_{430}\right) + \left(3\cdot\frac{5}{5}\cdot\boldsymbol{k}_{on}\boldsymbol{c}_{eff}\boldsymbol{y}_{450} - 2\cdot\boldsymbol{k}_{off}\boldsymbol{y}_{440}\right)$$

$$\frac{d\boldsymbol{y}_{431}}{dt} = \left(4\cdot\boldsymbol{k}_{on,comp}\boldsymbol{C}_{C}\boldsymbol{y}_{440} - 1\cdot\boldsymbol{k}_{off,comp}\boldsymbol{y}_{431}\right) - \left(3\cdot\boldsymbol{k}_{on,comp}\boldsymbol{C}_{C}\boldsymbol{y}_{431} - 2\cdot\boldsymbol{k}_{off,comp}\boldsymbol{y}_{422}\right) - \left(2\cdot\frac{3}{4}\cdot\boldsymbol{k}_{on}\boldsymbol{c}_{eff}\boldsymbol{y}_{431} - 3\cdot\boldsymbol{k}_{off}\boldsymbol{y}_{421}\right) + \left(3\cdot\frac{4}{5}\cdot\boldsymbol{k}_{on}\boldsymbol{c}_{eff}\boldsymbol{y}_{441} - 2\cdot\boldsymbol{k}_{off}\boldsymbol{y}_{431}\right)$$

$$\frac{d\boldsymbol{y}_{422}}{dt} = \left(3\cdot\boldsymbol{k}_{on,comp}\boldsymbol{C}_{C}\boldsymbol{y}_{431} - 2\cdot\boldsymbol{k}_{off,comp}\boldsymbol{y}_{422}\right) - \left(2\cdot\boldsymbol{k}_{on,comp}\boldsymbol{C}_{C}\boldsymbol{y}_{422} - 3\cdot\boldsymbol{k}_{off,comp}\boldsymbol{y}_{413}\right) - \left(2\cdot\frac{2}{4}\cdot\boldsymbol{k}_{on}\boldsymbol{c}_{eff}\boldsymbol{y}_{422} - 3\cdot\boldsymbol{k}_{off}\boldsymbol{y}_{412}\right) + \left(3\cdot\frac{3}{5}\cdot\boldsymbol{k}_{on}\boldsymbol{c}_{eff}\boldsymbol{y}_{432} - 2\cdot\boldsymbol{k}_{off}\boldsymbol{y}_{422}\right)$$

$$\frac{d\boldsymbol{y}_{413}}{dt} = \left(2\cdot\boldsymbol{k}_{on,comp}\boldsymbol{C}_{C}\boldsymbol{y}_{422} - 3\cdot\boldsymbol{k}_{off,comp}\boldsymbol{y}_{413}\right) - \left(1\cdot\boldsymbol{k}_{on,comp}\boldsymbol{C}_{C}\boldsymbol{y}_{413} - 4\cdot\boldsymbol{k}_{off,comp}\boldsymbol{y}_{404}\right) - \left(2\cdot\frac{1}{4}\cdot\boldsymbol{k}_{on}\boldsymbol{c}_{eff}\boldsymbol{y}_{413} - 3\cdot\boldsymbol{k}_{off}\boldsymbol{y}_{403}\right) + \left(3\cdot\frac{2}{5}\cdot\boldsymbol{k}_{on}\boldsymbol{c}_{eff}\boldsymbol{y}_{423} - 2\cdot\boldsymbol{k}_{off}\boldsymbol{y}_{413}\right)$$

$$\frac{d\boldsymbol{y}_{404}}{dt} = \left(1\cdot\boldsymbol{k}_{on,comp}\boldsymbol{C}_{C}\boldsymbol{y}_{413} - 4\cdot\boldsymbol{k}_{off,comp}\boldsymbol{y}_{404}\right) + \left(3\cdot\frac{1}{5}\cdot\boldsymbol{k}_{on}\boldsymbol{c}_{eff}\boldsymbol{y}_{414} - 2\cdot\boldsymbol{k}_{off}\boldsymbol{y}_{404}\right)$$

**3rd row (three gC1q heads bound):**

$$\frac{d\boldsymbol{y}_{430}}{dt} = -\left(3\cdot\boldsymbol{k}_{on,comp}\boldsymbol{C}_{C}\boldsymbol{y}_{430} - 1\cdot\boldsymbol{k}_{off,comp}\boldsymbol{y}_{421}\right) - \left(1\cdot\frac{3}{3}\cdot\boldsymbol{k}_{on}\boldsymbol{c}_{eff}\boldsymbol{y}_{430} - 4\cdot\boldsymbol{k}_{off}\boldsymbol{y}_{420}\right) + \left(2\cdot\frac{4}{4}\cdot\boldsymbol{k}_{on}\boldsymbol{c}_{eff}\boldsymbol{y}_{440} - 3\cdot\boldsymbol{k}_{off}\boldsymbol{y}_{430}\right)$$

$$\frac{d\boldsymbol{y}_{421}}{dt} = \left(3\cdot\boldsymbol{k}_{on,comp}\boldsymbol{C}_{C}\boldsymbol{y}_{430} - 1\cdot\boldsymbol{k}_{off,comp}\boldsymbol{y}_{421}\right) - \left(2\cdot\boldsymbol{k}_{on,comp}\boldsymbol{C}_{C}\boldsymbol{y}_{421} - 2\cdot\boldsymbol{k}_{off,comp}\boldsymbol{y}_{412}\right) - \left(1\cdot\frac{2}{3}\cdot\boldsymbol{k}_{on}\boldsymbol{c}_{eff}\boldsymbol{y}_{421} - 4\cdot\boldsymbol{k}_{off}\boldsymbol{y}_{411}\right) + \left(2\cdot\frac{3}{4}\cdot\boldsymbol{k}_{on}\boldsymbol{c}_{eff}\boldsymbol{y}_{431} - 3\cdot\boldsymbol{k}_{off}\boldsymbol{y}_{421}\right)$$

$$\frac{d\boldsymbol{y}_{412}}{dt} = \left(2\cdot\boldsymbol{k}_{on,comp}\boldsymbol{C}_{C}\boldsymbol{y}_{421} - 2\cdot\boldsymbol{k}_{off,comp}\boldsymbol{y}_{412}\right) - \left(1\cdot\boldsymbol{k}_{on,comp}\boldsymbol{C}_{C}\boldsymbol{y}_{412} - 3\cdot\boldsymbol{k}_{off,comp}\boldsymbol{y}_{403}\right) - \left(1\cdot\frac{1}{3}\cdot\boldsymbol{k}_{on}\boldsymbol{c}_{eff}\boldsymbol{y}_{412} - 4\cdot\boldsymbol{k}_{off}\boldsymbol{y}_{402}\right) + \left(2\cdot\frac{2}{4}\cdot\boldsymbol{k}_{on}\boldsymbol{c}_{eff}\boldsymbol{y}_{422} - 3\cdot\boldsymbol{k}_{off}\boldsymbol{y}_{412}\right)$$

$$\frac{d\boldsymbol{y}_{403}}{dt} = \left(1\cdot\boldsymbol{k}_{on,comp}\boldsymbol{C}_{C}\boldsymbol{y}_{412} - 3\cdot\boldsymbol{k}_{off,comp}\boldsymbol{y}_{403}\right) + \left(2\cdot\frac{1}{4}\cdot\boldsymbol{k}_{on}\boldsymbol{c}_{eff}\boldsymbol{y}_{413} - 3\cdot\boldsymbol{k}_{off}\boldsymbol{y}_{403}\right)$$

**4th row (four gC1q heads bound):**

$$\frac{d\boldsymbol{y}_{420}}{dt} = -\left(2\cdot\boldsymbol{k}_{on,comp}\boldsymbol{C}_{C}\boldsymbol{y}_{420} - 1\cdot\boldsymbol{k}_{off,comp}\boldsymbol{y}_{411}\right) + \left(1\cdot\frac{3}{3}\cdot\boldsymbol{k}_{on}\boldsymbol{c}_{eff}\boldsymbol{y}_{430} - 4\cdot\boldsymbol{k}_{off}\boldsymbol{y}_{420}\right)$$

$$\frac{d\boldsymbol{y}_{411}}{dt} = \left(2 \cdot \boldsymbol{k}_{on,comp}\boldsymbol{C}_C\boldsymbol{y}_{420} - 1 \cdot \boldsymbol{k}_{off,comp}\boldsymbol{y}_{411}\right) - \left(1 \cdot \boldsymbol{k}_{on,comp}\boldsymbol{C}_C\boldsymbol{y}_{411} - 2 \cdot \boldsymbol{k}_{off,comp}\boldsymbol{y}_{402}\right) + \left(1 \cdot \frac{2}{3} \cdot \boldsymbol{k}_{on}\boldsymbol{c}_{eff}\boldsymbol{y}_{421} - 4 \cdot \boldsymbol{k}_{off}\boldsymbol{y}_{411}\right)$$

$$\frac{d\boldsymbol{y}_{402}}{dt} = \left(1 \cdot \boldsymbol{k}_{on,comp}\boldsymbol{C}_C\boldsymbol{y}_{411} - 2 \cdot \boldsymbol{k}_{off,comp}\boldsymbol{y}_{402}\right) + \left(1 \cdot \frac{1}{3} \cdot \boldsymbol{k}_{on}\boldsymbol{c}_{eff}\boldsymbol{y}_{412} - 4 \cdot \boldsymbol{k}_{off}\boldsymbol{y}_{402}\right)$$

**Tetramers without C1/C1q:**

$$\frac{d\boldsymbol{y}_{4}}{dt} = -24 \cdot \boldsymbol{k}_{on}\boldsymbol{C}_{C1}\boldsymbol{y}_4 + \boldsymbol{k}_{off}(\boldsymbol{y}_{450} + \boldsymbol{y}_{441} + \boldsymbol{y}_{432} + \boldsymbol{y}_{423} + \boldsymbol{y}_{414} + \boldsymbol{y}_{405})$$

#### IV. C1/C1q binding to IgG Trimers (Fig. S5D):

**1st row (one gC1q head bound):**

$$\frac{d\boldsymbol{y}_{350}}{dt} = -\left(5 \cdot \boldsymbol{k}_{on,comp}\boldsymbol{C}_C\boldsymbol{y}_{350} - 1 \cdot \boldsymbol{k}_{off,comp}\boldsymbol{y}_{341}\right) - \left(2 \cdot \frac{5}{5} \cdot \boldsymbol{k}_{on}\boldsymbol{c}_{eff}\boldsymbol{y}_{350} - 2 \cdot \boldsymbol{k}_{off}\boldsymbol{y}_{340}\right) + \left(18 \cdot \boldsymbol{k}_{on}\boldsymbol{C}_{C1}\boldsymbol{y}_3 - \boldsymbol{k}_{off}\boldsymbol{y}_{350}\right)$$

$$\frac{d\boldsymbol{y}_{341}}{dt} = \left(5 \cdot \boldsymbol{k}_{on,comp}\boldsymbol{C}_C\boldsymbol{y}_{350} - 1 \cdot \boldsymbol{k}_{off,comp}\boldsymbol{y}_{341}\right) - \left(4 \cdot \boldsymbol{k}_{on,comp}\boldsymbol{C}_C\boldsymbol{y}_{341} - 2 \cdot \boldsymbol{k}_{off,comp}\boldsymbol{y}_{332}\right) - \left(2 \cdot \frac{4}{5} \cdot \boldsymbol{k}_{on}\boldsymbol{c}_{eff}\boldsymbol{y}_{341} - 2 \cdot \boldsymbol{k}_{off}\boldsymbol{y}_{331}\right) + \left( -\boldsymbol{k}_{off}\boldsymbol{y}_{341}\right)$$

$$\frac{d\boldsymbol{y}_{332}}{dt} = \left(4 \cdot \boldsymbol{k}_{on,comp}\boldsymbol{C}_C\boldsymbol{y}_{341} - 2 \cdot \boldsymbol{k}_{off,comp}\boldsymbol{y}_{332}\right) - \left(3 \cdot \boldsymbol{k}_{on,comp}\boldsymbol{C}_C\boldsymbol{y}_{332} - 3 \cdot \boldsymbol{k}_{off,comp}\boldsymbol{y}_{323}\right) - \left(2 \cdot \frac{3}{5} \cdot \boldsymbol{k}_{on}\boldsymbol{c}_{eff}\boldsymbol{y}_{332} - 2 \cdot \boldsymbol{k}_{off}\boldsymbol{y}_{322}\right) + \left( -\boldsymbol{k}_{off}\boldsymbol{y}_{332}\right)$$

$$\frac{d\boldsymbol{y}_{323}}{dt} = \left(3 \cdot \boldsymbol{k}_{on,comp}\boldsymbol{C}_C\boldsymbol{y}_{332} - 3 \cdot \boldsymbol{k}_{off,comp}\boldsymbol{y}_{323}\right) - \left(2 \cdot \boldsymbol{k}_{on,comp}\boldsymbol{C}_C\boldsymbol{y}_{323} - 4 \cdot \boldsymbol{k}_{off,comp}\boldsymbol{y}_{314}\right) - \left(2 \cdot \frac{2}{5} \cdot \boldsymbol{k}_{on}\boldsymbol{c}_{eff}\boldsymbol{y}_{323} - 2 \cdot \boldsymbol{k}_{off}\boldsymbol{y}_{313}\right) + \left( -\boldsymbol{k}_{off}\boldsymbol{y}_{323}\right)$$

$$\frac{d\boldsymbol{y}_{314}}{dt} = \left(2 \cdot \boldsymbol{k}_{on,comp}\boldsymbol{C}_C\boldsymbol{y}_{323} - 4 \cdot \boldsymbol{k}_{off,comp}\boldsymbol{y}_{314}\right) - \left(1 \cdot \boldsymbol{k}_{on,comp}\boldsymbol{C}_C\boldsymbol{y}_{314} - 5 \cdot \boldsymbol{k}_{off,comp}\boldsymbol{y}_{305}\right) - \left(2 \cdot \frac{1}{5} \cdot \boldsymbol{k}_{on}\boldsymbol{c}_{eff}\boldsymbol{y}_{314} - 2 \cdot \boldsymbol{k}_{off}\boldsymbol{y}_{304}\right) + \left( -\boldsymbol{k}_{off}\boldsymbol{y}_{314}\right)$$

$$\frac{d\boldsymbol{y}_{305}}{dt} = \left(1 \cdot \boldsymbol{k}_{on,comp}\boldsymbol{C}_C\boldsymbol{y}_{314} - 5 \cdot \boldsymbol{k}_{off,comp}\boldsymbol{y}_{305}\right) + \left( -\boldsymbol{k}_{off}\boldsymbol{y}_{305}\right)$$

**2nd row (two gC1q heads bound):**

$$\frac{d\boldsymbol{y}_{340}}{dt} = -\left(4 \cdot \boldsymbol{k}_{on,comp}\boldsymbol{C}_C\boldsymbol{y}_{340} - 1 \cdot \boldsymbol{k}_{off,comp}\boldsymbol{y}_{331}\right) - \left(1 \cdot \frac{4}{4} \cdot \boldsymbol{k}_{on}\boldsymbol{c}_{eff}\boldsymbol{y}_{340} - 3 \cdot \boldsymbol{k}_{off}\boldsymbol{y}_{330}\right) + \left(2 \cdot \frac{5}{5} \cdot \boldsymbol{k}_{on}\boldsymbol{c}_{eff}\boldsymbol{y}_{350} - 2 \cdot \boldsymbol{k}_{off}\boldsymbol{y}_{340}\right)$$

$$\frac{d\boldsymbol{y}_{331}}{dt} = \left(4 \cdot \boldsymbol{k}_{on,comp}\boldsymbol{C}_C\boldsymbol{y}_{340} - 1 \cdot \boldsymbol{k}_{off,comp}\boldsymbol{y}_{331}\right) - \left(3 \cdot \boldsymbol{k}_{on,comp}\boldsymbol{C}_C\boldsymbol{y}_{331} - 2 \cdot \boldsymbol{k}_{off,comp}\boldsymbol{y}_{322}\right) - \left(1 \cdot \frac{3}{4} \cdot \boldsymbol{k}_{on}\boldsymbol{c}_{eff}\boldsymbol{y}_{331} - 3 \cdot \boldsymbol{k}_{off}\boldsymbol{y}_{321}\right) + \left(2 \cdot \frac{4}{5} \cdot \boldsymbol{k}_{on}\boldsymbol{c}_{eff}\boldsymbol{y}_{341} - 2 \cdot \boldsymbol{k}_{off}\boldsymbol{y}_{331}\right)$$

$$\frac{d\boldsymbol{y}_{322}}{dt} = \left(3 \cdot \boldsymbol{k}_{on,comp}\boldsymbol{C}_C\boldsymbol{y}_{331} - 2 \cdot \boldsymbol{k}_{off,comp}\boldsymbol{y}_{322}\right) - \left(2 \cdot \boldsymbol{k}_{on,comp}\boldsymbol{C}_C\boldsymbol{y}_{322} - 3 \cdot \boldsymbol{k}_{off,comp}\boldsymbol{y}_{313}\right) - \left(1 \cdot \frac{2}{4} \cdot \boldsymbol{k}_{on}\boldsymbol{c}_{eff}\boldsymbol{y}_{322} - 3 \cdot \boldsymbol{k}_{off}\boldsymbol{y}_{312}\right) + \left(2 \cdot \frac{3}{5} \cdot \boldsymbol{k}_{on}\boldsymbol{c}_{eff}\boldsymbol{y}_{332} - 2 \cdot \boldsymbol{k}_{off}\boldsymbol{y}_{322}\right)$$

$$\frac{d\boldsymbol{y}_{313}}{dt} = \left(2 \cdot \boldsymbol{k}_{on,comp}\boldsymbol{C}_C\boldsymbol{y}_{322} - 3 \cdot \boldsymbol{k}_{off,comp}\boldsymbol{y}_{313}\right) - \left(1 \cdot \boldsymbol{k}_{on,comp}\boldsymbol{C}_C\boldsymbol{y}_{313} - 4 \cdot \boldsymbol{k}_{off,comp}\boldsymbol{y}_{304}\right) - \left(1 \cdot \frac{1}{4} \cdot \boldsymbol{k}_{on}\boldsymbol{c}_{eff}\boldsymbol{y}_{313} - 3 \cdot \boldsymbol{k}_{off}\boldsymbol{y}_{303}\right) + \left(2 \cdot \frac{2}{5} \cdot \boldsymbol{k}_{on}\boldsymbol{c}_{eff}\boldsymbol{y}_{323} - 2 \cdot \boldsymbol{k}_{off}\boldsymbol{y}_{313}\right)$$

$$\frac{d\boldsymbol{y}_{304}}{dt} = \left(1 \cdot \boldsymbol{k}_{on,comp}\boldsymbol{C}_C\boldsymbol{y}_{313} - 4 \cdot \boldsymbol{k}_{off,comp}\boldsymbol{y}_{304}\right) + \left(2 \cdot \frac{1}{5} \cdot \boldsymbol{k}_{on}\boldsymbol{c}_{eff}\boldsymbol{y}_{314} - 2 \cdot \boldsymbol{k}_{off}\boldsymbol{y}_{304}\right)$$

**3rd row (three gC1q heads bound):**

$$\frac{d\boldsymbol{y}_{330}}{dt} = -\left(3\cdot\boldsymbol{k}_{on,comp}\boldsymbol{C}_C\boldsymbol{y}_{330} - 1\cdot\boldsymbol{k}_{off,comp}\boldsymbol{y}_{321}\right) + \left(1\cdot\frac{4}{4}\cdot\boldsymbol{k}_{on}\boldsymbol{c}_{eff}\boldsymbol{y}_{340} - 3\cdot\boldsymbol{k}_{off}\boldsymbol{y}_{330}\right)$$

$$\frac{d\boldsymbol{y}_{321}}{dt} = \left(3\cdot\boldsymbol{k}_{on,comp}\boldsymbol{C}_C\boldsymbol{y}_{330} - 1\cdot\boldsymbol{k}_{off,comp}\boldsymbol{y}_{321}\right) - \left(2\cdot\boldsymbol{k}_{on,comp}\boldsymbol{C}_C\boldsymbol{y}_{321} - 2\cdot\boldsymbol{k}_{off,comp}\boldsymbol{y}_{312}\right) + \left(1\cdot\frac{3}{4}\cdot\boldsymbol{k}_{on}\boldsymbol{c}_{eff}\boldsymbol{y}_{331} - 3\cdot\boldsymbol{k}_{off}\boldsymbol{y}_{321}\right)$$

$$\frac{d\boldsymbol{y}_{312}}{dt} = \left(2\cdot\boldsymbol{k}_{on,comp}\boldsymbol{C}_C\boldsymbol{y}_{321} - 2\cdot\boldsymbol{k}_{off,comp}\boldsymbol{y}_{312}\right) - \left(1\cdot\boldsymbol{k}_{on,comp}\boldsymbol{C}_C\boldsymbol{y}_{312} - 3\cdot\boldsymbol{k}_{off,comp}\boldsymbol{y}_{303}\right) + \left(1\cdot\frac{2}{4}\cdot\boldsymbol{k}_{on}\boldsymbol{c}_{eff}\boldsymbol{y}_{322} - 3\cdot\boldsymbol{k}_{off}\boldsymbol{y}_{312}\right)$$

$$\frac{d\boldsymbol{y}_{303}}{dt} = \left(1\cdot\boldsymbol{k}_{on,comp}\boldsymbol{C}_C\boldsymbol{y}_{312} - 3\cdot\boldsymbol{k}_{off,comp}\boldsymbol{y}_{303}\right) + \left(1\cdot\frac{1}{4}\cdot\boldsymbol{k}_{on}\boldsymbol{c}_{eff}\boldsymbol{y}_{313} - 3\cdot\boldsymbol{k}_{off}\boldsymbol{y}_{303}\right)$$

**Trimers without C1/C1q:**

$$\frac{d\boldsymbol{y}_{3}}{dt} = -18\cdot\boldsymbol{k}_{on}\boldsymbol{C}_{C1}\boldsymbol{y}_{3} + \boldsymbol{k}_{off}(\boldsymbol{y}_{350} + \boldsymbol{y}_{341} + \boldsymbol{y}_{332} + \boldsymbol{y}_{323} + \boldsymbol{y}_{314} + \boldsymbol{y}_{305})$$

#### V. C1/C1q binding to IgG Dimers (Fig. S5E):

**1st row (one gC1q head bound):**

$$\frac{d\boldsymbol{y}_{250}}{dt} = -\left(5\cdot\boldsymbol{k}_{on,comp}\boldsymbol{C}_C\boldsymbol{y}_{250} - 1\cdot\boldsymbol{k}_{off,comp}\boldsymbol{y}_{241}\right) - \left(1\cdot\frac{5}{5}\cdot\boldsymbol{k}_{on}\boldsymbol{c}_{eff}\boldsymbol{y}_{250} - 2\cdot\boldsymbol{k}_{off}\boldsymbol{y}_{240}\right) + \left(12\cdot\boldsymbol{k}_{on}\boldsymbol{C}_{C1}\boldsymbol{y}_{2} - \boldsymbol{k}_{off}\boldsymbol{y}_{250}\right)$$

$$\frac{d\boldsymbol{y}_{241}}{dt} = \left(5\cdot\boldsymbol{k}_{on,comp}\boldsymbol{C}_C\boldsymbol{y}_{250} - 1\cdot\boldsymbol{k}_{off,comp}\boldsymbol{y}_{241}\right) - \left(4\cdot\boldsymbol{k}_{on,comp}\boldsymbol{C}_C\boldsymbol{y}_{241} - 2\cdot\boldsymbol{k}_{off,comp}\boldsymbol{y}_{232}\right) - \left(1\cdot\frac{4}{5}\cdot\boldsymbol{k}_{on}\boldsymbol{c}_{eff}\boldsymbol{y}_{241} - 2\cdot\boldsymbol{k}_{off}\boldsymbol{y}_{231}\right) + \left(-\boldsymbol{k}_{off}\boldsymbol{y}_{241}\right)$$

$$\frac{d\boldsymbol{y}_{232}}{dt} = \left(4\cdot\boldsymbol{k}_{on,comp}\boldsymbol{C}_C\boldsymbol{y}_{241} - 2\cdot\boldsymbol{k}_{off,comp}\boldsymbol{y}_{232}\right) - \left(3\cdot\boldsymbol{k}_{on,comp}\boldsymbol{C}_C\boldsymbol{y}_{232} - 3\cdot\boldsymbol{k}_{off,comp}\boldsymbol{y}_{223}\right) - \left(1\cdot\frac{3}{5}\cdot\boldsymbol{k}_{on}\boldsymbol{c}_{eff}\boldsymbol{y}_{232} - 2\cdot\boldsymbol{k}_{off}\boldsymbol{y}_{222}\right) + \left(-\boldsymbol{k}_{off}\boldsymbol{y}_{232}\right)$$

$$\frac{d\boldsymbol{y}_{223}}{dt} = \left(3\cdot\boldsymbol{k}_{on,comp}\boldsymbol{C}_C\boldsymbol{y}_{232} - 3\cdot\boldsymbol{k}_{off,comp}\boldsymbol{y}_{223}\right) - \left(2\cdot\boldsymbol{k}_{on,comp}\boldsymbol{C}_C\boldsymbol{y}_{223} - 4\cdot\boldsymbol{k}_{off,comp}\boldsymbol{y}_{214}\right) - \left(1\cdot\frac{2}{5}\cdot\boldsymbol{k}_{on}\boldsymbol{c}_{eff}\boldsymbol{y}_{223} - 2\cdot\boldsymbol{k}_{off}\boldsymbol{y}_{213}\right) + \left(-\boldsymbol{k}_{off}\boldsymbol{y}_{223}\right)$$

$$\frac{d\boldsymbol{y}_{214}}{dt} = \left(2\cdot\boldsymbol{k}_{on,comp}\boldsymbol{C}_C\boldsymbol{y}_{223} - 4\cdot\boldsymbol{k}_{off,comp}\boldsymbol{y}_{214}\right) - \left(1\cdot\boldsymbol{k}_{on,comp}\boldsymbol{C}_C\boldsymbol{y}_{214} - 5\cdot\boldsymbol{k}_{off,comp}\boldsymbol{y}_{205}\right) - \left(1\cdot\frac{1}{5}\cdot\boldsymbol{k}_{on}\boldsymbol{c}_{eff}\boldsymbol{y}_{214} - 2\cdot\boldsymbol{k}_{off}\boldsymbol{y}_{204}\right) + \left(-\boldsymbol{k}_{off}\boldsymbol{y}_{214}\right)$$

$$\frac{d\boldsymbol{y}_{205}}{dt} = \left(1\cdot\boldsymbol{k}_{on,comp}\boldsymbol{C}_C\boldsymbol{y}_{214} - 5\cdot\boldsymbol{k}_{off,comp}\boldsymbol{y}_{205}\right) + \left(-\boldsymbol{k}_{off}\boldsymbol{y}_{205}\right)$$

**2nd row (two gC1q heads bound):**

$$\frac{d\boldsymbol{y}_{240}}{dt} = -\left(4\cdot\boldsymbol{k}_{on,comp}\boldsymbol{C}_C\boldsymbol{y}_{240} - 1\cdot\boldsymbol{k}_{off,comp}\boldsymbol{y}_{31}\right) - + \left(1\cdot\frac{5}{5}\cdot\boldsymbol{k}_{on}\boldsymbol{c}_{eff}\boldsymbol{y}_{250} - 2\cdot\boldsymbol{k}_{off}\boldsymbol{y}_{240}\right)$$

$$\frac{d\boldsymbol{y}_{231}}{dt} = \left(4\cdot\boldsymbol{k}_{on,comp}\boldsymbol{C}_C\boldsymbol{y}_{240} - 1\cdot\boldsymbol{k}_{off,comp}\boldsymbol{y}_{231}\right) - \left(3\cdot\boldsymbol{k}_{on,comp}\boldsymbol{C}_C\boldsymbol{y}_{231} - 2\cdot\boldsymbol{k}_{off,comp}\boldsymbol{y}_{222}\right) - + \left(1\cdot\frac{4}{5}\cdot\boldsymbol{k}_{on}\boldsymbol{c}_{eff}\boldsymbol{y}_{241} - 2\cdot\boldsymbol{k}_{off}\boldsymbol{y}_{231}\right)$$

$$\frac{d\boldsymbol{y}_{222}}{dt} = \left(3\cdot\boldsymbol{k}_{on,comp}\boldsymbol{C}_C\boldsymbol{y}_{231} - 2\cdot\boldsymbol{k}_{off,comp}\boldsymbol{y}_{222}\right) - \left(2\cdot\boldsymbol{k}_{on,comp}\boldsymbol{C}_C\boldsymbol{y}_{222} - 3\cdot\boldsymbol{k}_{off,comp}\boldsymbol{y}_{213}\right) - + \left(1\cdot\frac{3}{5}\cdot\boldsymbol{k}_{on}\boldsymbol{c}_{eff}\boldsymbol{y}_{232} - 2\cdot\boldsymbol{k}_{off}\boldsymbol{y}_{222}\right)$$

$$\frac{d\boldsymbol{y}_{213}}{dt} = \left(2\cdot\boldsymbol{k}_{on,comp}\boldsymbol{C}_C\boldsymbol{y}_{222} - 3\cdot\boldsymbol{k}_{off,comp}\boldsymbol{y}_{213}\right) - \left(1\cdot\boldsymbol{k}_{on,comp}\boldsymbol{C}_C\boldsymbol{y}_{213} - 4\cdot\boldsymbol{k}_{off,comp}\boldsymbol{y}_{204}\right) - + \left(1\cdot\frac{2}{5}\cdot\boldsymbol{k}_{on}\boldsymbol{c}_{eff}\boldsymbol{y}_{223} - 2\cdot\boldsymbol{k}_{off}\boldsymbol{y}_{213}\right)$$

$$\frac{d\boldsymbol{y}_{204}}{dt} = \left(1 \cdot \boldsymbol{k}_{on,comp}\boldsymbol{C}_C\boldsymbol{y}_{213} - 4 \cdot \boldsymbol{k}_{off,comp}\boldsymbol{y}_{204}\right) + \left(1 \cdot \frac{1}{5} \cdot \boldsymbol{k}_{on}\boldsymbol{c}_{eff}\boldsymbol{y}_{214} - 2 \cdot \boldsymbol{k}_{off}\boldsymbol{y}_{204}\right)$$

**Dimers without C1/C1q:**

$$\frac{d\boldsymbol{y}_{2}}{dt} = -12 \cdot \boldsymbol{k}_{on}\boldsymbol{C}_{C1}\boldsymbol{y}_2 + \boldsymbol{k}_{off}(\boldsymbol{y}_{250} + \boldsymbol{y}_{241} + \boldsymbol{y}_{232} + \boldsymbol{y}_{223} + \boldsymbol{y}_{214} + \boldsymbol{y}_{205})$$

#### VI. C1/C1q binding to IgG Monomers (Fig. S5F):

**1st row (one gC1q head bound):**

$$\frac{d\boldsymbol{y}_{150}}{dt} = -\left(5 \cdot \boldsymbol{k}_{on,comp}\boldsymbol{C}_C\boldsymbol{y}_{150} - 1 \cdot \boldsymbol{k}_{off,comp}\boldsymbol{y}_{141}\right) - +(6 \cdot \boldsymbol{k}_{on}\boldsymbol{C}_{C1}\boldsymbol{y}_1 - \boldsymbol{k}_{off}\boldsymbol{y}_{150})$$

$$\frac{d\boldsymbol{y}_{141}}{dt} = \left(5 \cdot \boldsymbol{k}_{on,comp}\boldsymbol{C}_C\boldsymbol{y}_{150} - 1 \cdot \boldsymbol{k}_{off,comp}\boldsymbol{y}_{141}\right) - \left(4 \cdot \boldsymbol{k}_{on,comp}\boldsymbol{C}_C\boldsymbol{y}_{141} - 2 \cdot \boldsymbol{k}_{off,comp}\boldsymbol{y}_{132}\right) - +( \quad -\boldsymbol{k}_{off}\boldsymbol{y}_{141})$$

$$\frac{d\boldsymbol{y}_{132}}{dt} = \left(4 \cdot \boldsymbol{k}_{on,comp}\boldsymbol{C}_C\boldsymbol{y}_{141} - 2 \cdot \boldsymbol{k}_{off,comp}\boldsymbol{y}_{132}\right) - \left(3 \cdot \boldsymbol{k}_{on,comp}\boldsymbol{C}_C\boldsymbol{y}_{132} - 3 \cdot \boldsymbol{k}_{off,comp}\boldsymbol{y}_{123}\right) - +( \quad -\boldsymbol{k}_{off}\boldsymbol{y}_{132})$$

$$\frac{d\boldsymbol{y}_{123}}{dt} = \left(3 \cdot \boldsymbol{k}_{on,comp}\boldsymbol{C}_C\boldsymbol{y}_{132} - 3 \cdot \boldsymbol{k}_{off,comp}\boldsymbol{y}_{123}\right) - \left(2 \cdot \boldsymbol{k}_{on,comp}\boldsymbol{C}_C\boldsymbol{y}_{123} - 4 \cdot \boldsymbol{k}_{off,comp}\boldsymbol{y}_{114}\right) - +( \quad -\boldsymbol{k}_{off}\boldsymbol{y}_{123})$$

$$\frac{d\boldsymbol{y}_{114}}{dt} = \left(2 \cdot \boldsymbol{k}_{on,comp}\boldsymbol{C}_C\boldsymbol{y}_{123} - 4 \cdot \boldsymbol{k}_{off,comp}\boldsymbol{y}_{114}\right) - \left(1 \cdot \boldsymbol{k}_{on,comp}\boldsymbol{C}_C\boldsymbol{y}_{114} - 5 \cdot \boldsymbol{k}_{off,comp}\boldsymbol{y}_{105}\right) - +( \quad -\boldsymbol{k}_{off}\boldsymbol{y}_{114})$$

$$\frac{d\boldsymbol{y}_{105}}{dt} = \left(1 \cdot \boldsymbol{k}_{on,comp}\boldsymbol{C}_C\boldsymbol{y}_{114} - 5 \cdot \boldsymbol{k}_{off,comp}\boldsymbol{y}_{105}\right) + ( \quad -\boldsymbol{k}_{off}\boldsymbol{y}_{105})$$

**Monomers without C1/C1q:**

$$\frac{d\boldsymbol{y}_{1}}{dt} = -6 \cdot \boldsymbol{k}_{on}\boldsymbol{C}_{C1}\boldsymbol{y}_1 + \boldsymbol{k}_{off}(\boldsymbol{y}_{150} + \boldsymbol{y}_{141} + \boldsymbol{y}_{132} + \boldsymbol{y}_{123} + \boldsymbol{y}_{114} + \boldsymbol{y}_{105})$$

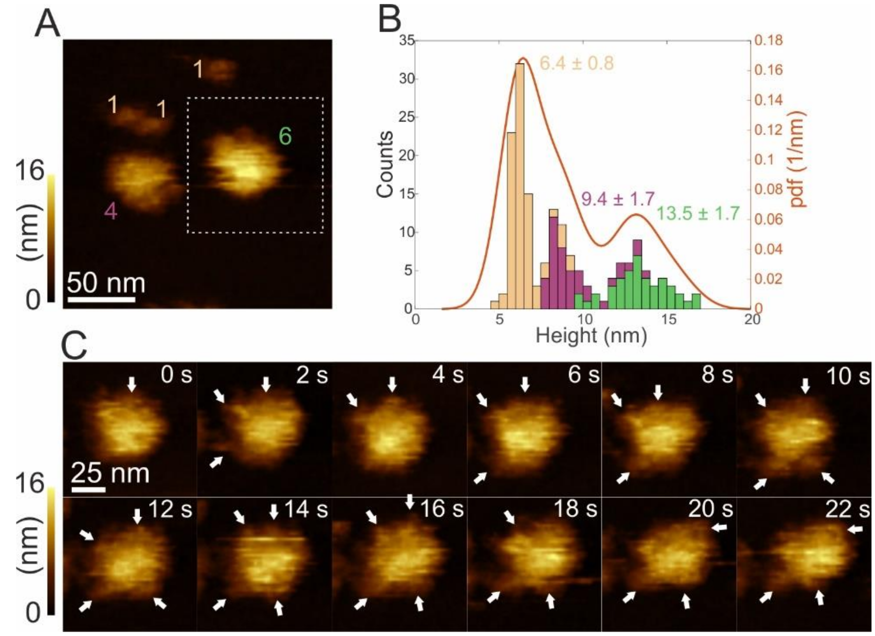


**Figure S1.** Structural comparison of IgG3 and IgG1 hexamers based on higher-resolution HS-AFM images. **(A)** First frame of HS-AFM movie S2 of one IgG3 hexamer, one tetramer and some monomers bound to a DNP-SLB. **(B)** Height histogram generated from height over time recordings of the oligomers in movie S2. Contributions of the respective oligomers are color coded according to (A). Numbers correspond to means ± s.d. over all image frames and individual particles, respectively. **(C)** High resolution images of an individual IgG3 hexamer (dashed area from (A)). Additional smaller structures surrounding the central Fc platform are indicated by arrows.

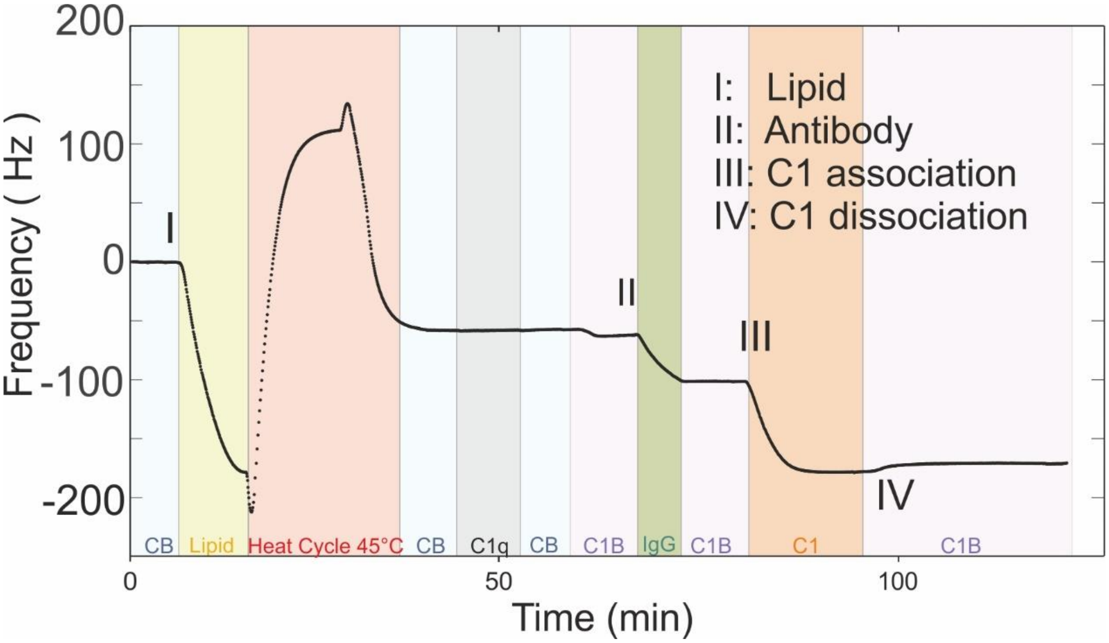


**Figure S2.** Typical QCM sensorgram of C1/C1q binding experiments. After equilibration with running buffer (CB), a lipid suspension (200 µg/ml) of DNP-labeled liposomes is injected into the QCM liquid cell (I) until saturation is reached, followed by a heat cycle to a maximum temperature of 45° C to facilitate liposome spreading and fusion on the $SiO_2$-coated QCM crystal forming a dense DNP-SLB. After removing excess lipids by flushing the DNP-SLB with CB, C1q is added to the running buffer at 15 nM to check for lipid membrane integrity (C1q would strongly associate with the bare $SiO_2$ surface when not covered by a lipid membrane). When no C1q binding was observed, the running buffer was changed from CB to C1B, and an anti-DNP IgG suspension (33 nM) was introduced (II) until the desired antibody density was reached. After removal of solution-phase IgGs through flushing with C1B, C1 or C1q was added at 15 nM until equilibrium was reached, after which a dissociation phase in C1B was added (IV). Data shown corresponds to the third overtone $f_3$.

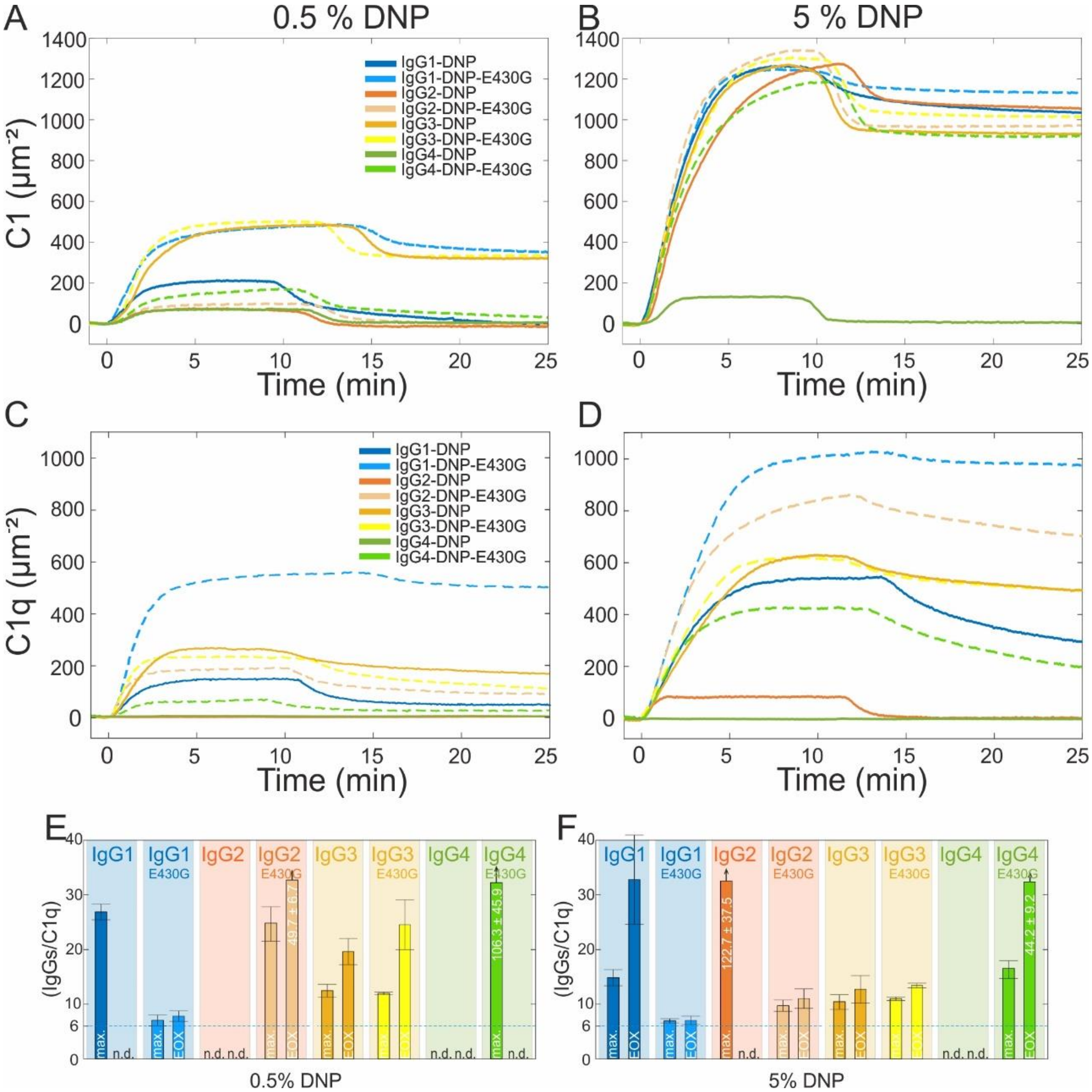


**Figure S3.** QCM sensorgrams of C1/C1q binding to IgG subclass variants and impact of antigen surface density on C1q recruitment efficiencies. **(A-B)** C1 binding to IgG subclass variants bound to 0.5 mol% DNP-SLBs (A) and 5 mol% DNP-SLBs (B). **(C-D)** C1q binding to IgG subclass variants bound to 0.5 mol% DNP-SLBs (C) and 5 mol% DNP-SLBs (D). **(E)** C1q recruitment efficiencies obtained for a medium antigen density of 0.5 mol % DNP-labelled lipids in the DNP-SLB. **(F)** C1q recruitment efficiencies of IgG1 and IgG3 variants obtained for a high antigen density of 5 mol% DNP-labelled lipids in the DNP-SLB. Depicted recruitment efficiencies are means ± s.d..

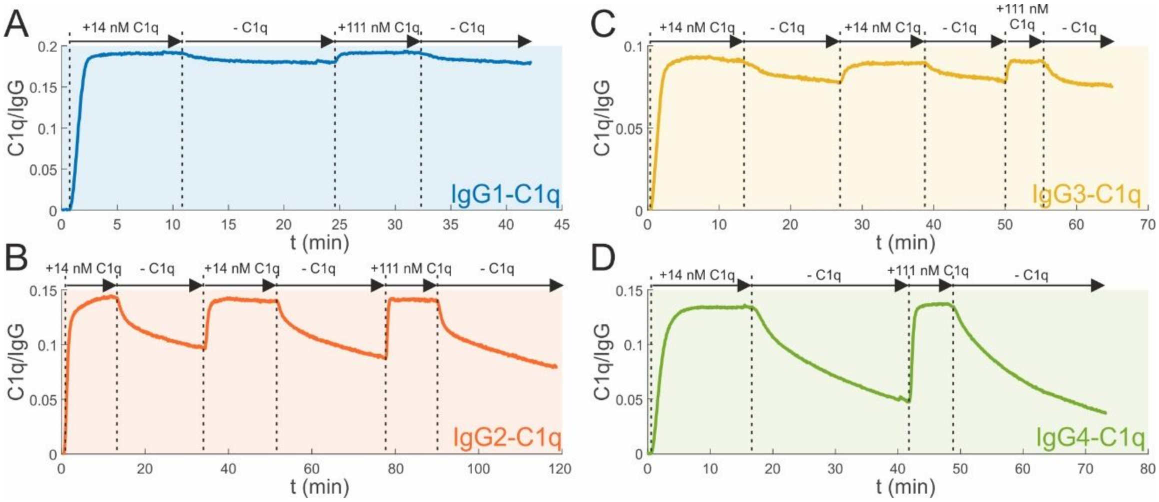


**Figure S4.** C1q binding to IgG-DNP-RGY opsonized DNP-SLBs saturates at the used C1q concentrations. **(A)** IgG1-DNP-RGY, **(B)** IgG2-DNP-RGY, **(C)** IgG3-DNP-RGY, **(D)** and IgG4-DNP-RGY. No significant additional binding was observed when 8-fold increasing the C1q concentration in the running buffer.

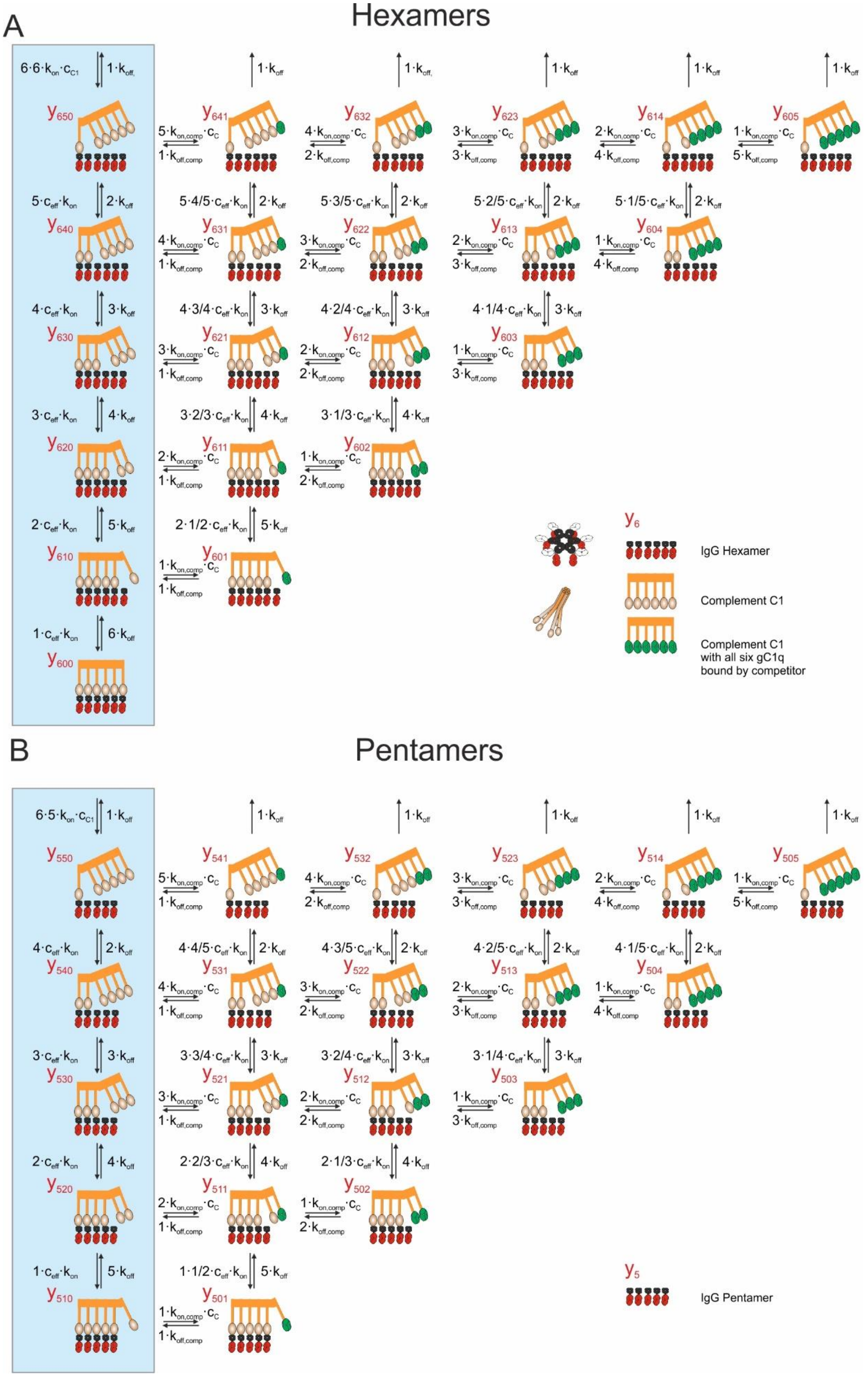
A
Hexamers
$y_6$
IgG Hexamer
Complement C1
Complement C1
with all six gC1q
bound by competitor
B
Pentamers
$y_5$
IgG Pentamer

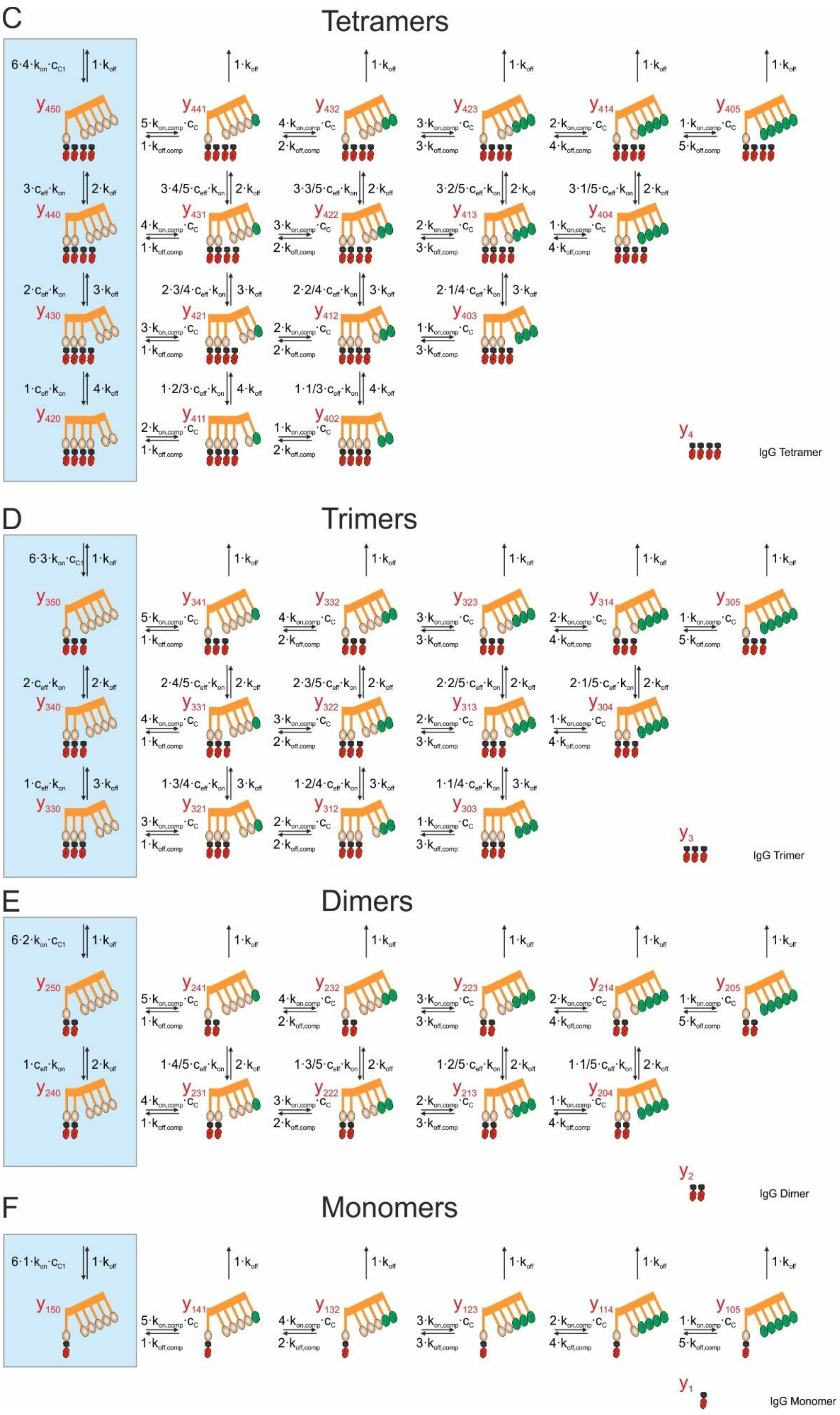
C
Tetramers
IgG Tetramer
D
Trimers
IgG Trimer
E
Dimers
IgG Dimer
F
Monomers
IgG Monomer

**Figure S5.** Mechanistic model of C1/C1q bound to IgG monomers-hexamers **(A-F)** in the absence (blue background) / presence of a competitor for gC1q. The interaction between the competitor and solution phase C1/C1q is not considered since the latter is already removed from solution when the competitor is introduced in our QCM experiments (Fig. 6).

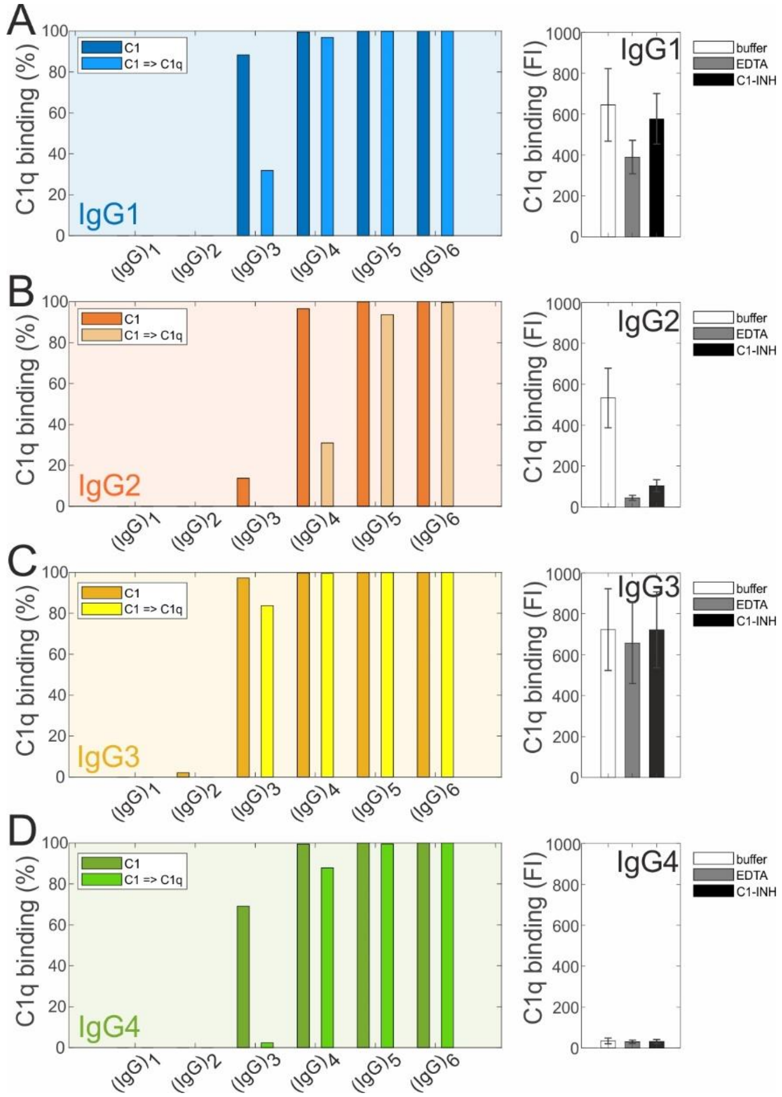


**Figure S6.** Model predictions for the removal of $C1r_2s_2$ from IgG-oligomer bound C1 by C1-INH or EDTA (right panels, data taken from (1)) on the stability of the remaining C1q – IgG oligomer complexes compared to the stability of the respective C1- IgG oligomer complexes. Following the experimental protocol given in (1), we simulated the binding of 3 nM C1 to IgG1-4 oligomers (incubation time: 30 min), after which the effective concentration $c_{eff}$ (cf. Fig. 6B, Table 2) was either switched from 1.5 to 0.4 mM (light bars), or left at 1.5 mM (dark bars) mimicking the effect of $C1r_2s_2$ removal/retention. Bars represent the percentage of C1q molecules remaining bound to the respective IgG oligomers 60 min after the end of the incubation time, when in the experiment C1q binding was detected in a flow cytometer. At this point, both C1 and C1q are entirely dissociated from IgG1-4 monomers and dimers (except some residual ~2 % C1 bound to IgG3 dimers), while removal of $C1r_2s_2$ from C1 affects C1q binding to higher IgG oligomers (> dimers) differently. **(A)** For IgG1, removal of $C1r_2s_2$ results in a reduction of C1q binding to IgG3 trimers from ~ 90 % to 30 %, while IgG1 tetramers to hexamers are vastly unaffected. The experimentally observed reduction in C1q binding (right panel) thus likely reflects the dissociation of C1q from IgG1 trimers. **(B)** In case of IgG2, only ~ 15 % of IgG2 trimers are predicted to be occupied by C1, which entirely dissociate when $C1r_2s_2$ is removed. C1q binding to IgG2 tetramers reduces to ~ 30 % upon removal of the proteases as compared to almost 100 % occupation by C1, while IgG2 pentamers and hexamers would both strongly bind C1 and C1q. This suggests that the strong reduction in C1q binding observed experimentally (right panel) is caused by the dissociation from IgG2 trimers and tetramers, while only a very small amount (if any) IgG2 pentamers and hexamers were present on the antigenic surface. Consequently, the C3b deposition observed for IgG2 (which was comparable to IgG1 and IgG3 (1)) likely largely originates from C1 activation by IgG2 tetramers. **(C)** $C1r_2s_2$ removal leads to only minor dissociation of C1q from IgG3 trimers and does not affect binding to tetramers, pentamers and hexamers, which again correctly predicts the experimental observations (right panel). **(D)** While our model predicts strong C1 and C1q binding to hypothetical IgG4 tetramers-hexamers and a strong reduction of C1q binding for the removal of $C1r_2s_2$ from C1 bound to IgG4 trimers, neither C1/C1q binding (right panel) nor C3b deposition was experimentally observed for IgG4 (1), suggesting that IgG4 solely was present in form of monomers or dimers on the antigenic surface and was thus not detected, similar to what we have observed in our QCM experiments (Fig. 5).

**Movie S1 (separate file).** HS-AFM movie of two IgG3-DNP hexamers and four IgG3-DNP monomers bound to a DNP-SLB recorded at a scan speed of 2 s/frame. Scan size: 200 x 200 $nm^2$ (100 x 100 pixel). Color scale range: 0- 15 nm.

**Movie S2 (separate file).** HS-AFM movie of an IgG3-DNP hexamer, an IgG3-DNP tetramer and IgG3-DNP monomers bound to a DNP-SLB recorded at a scan speed of 2 s/frame. Scan size: 200 x 200 $nm^2$ (100 x 100 pixel). Color scale range: 0- 16 nm.

**Movie S3 (separate file).** HS-AFM movie of an IgG1-DNP hexamer and an IgG1-DNP monomer bound to a DNP-SLB recorded at a scan speed of 2 s/frame. Scan size: 200 x 200 $nm^2$ (100 x 100 pixel). Color scale range: 0- 13 nm.

**Movie S4 (separate file).** HS-AFM movie of an IgG3-DNP hexamer bound to a DNP-SLB recorded at a scan speed of 2 s/frame. Scan size: 100 x 100 $nm^2$ (100 x 100 pixel). Color scale range: 0- 15 nm.
